# miR-100-5p modulates postprandial triglyceride response by targeting PCSK9

**DOI:** 10.64898/2026.03.26.713909

**Authors:** Amandine Vanduyse, Alexandre Motte, Carolina Neves, Rita Daclat, Sophie Galier, Olivier Bluteau, Clément Materne, Eric Frisdal, Hervé Durand, Philippe Giral, Joe-Elie Salem, Jean-Marc Lacorte, RESIST-PP Consortium, Cedric Le May, Wilfried Le Goff, Philippe Lesnik, Maryse Guerin

## Abstract

**Background:** Elevated postprandial hypertriglyceridemia (PP-HTG) is a significant risk factor for development of cardiovascular diseases, however, the mechanisms underlying its exaggerated rise remains poorly understood. MicroRNAs (miRs) are known to be implicated in the regulation of lipid metabolism, thus identifying them as potential key players. We presently investigated whether miRs may control postprandial triglyceride (PP-TG) response.

**Methods:** Postprandial changes in circulating miR expression as a function of the degree of postprandial TG response were evaluated in non-dyslipidemic healthy subjects (n=32). The impact of miR-100-5p on hepatic gene expression was evaluated in differentiated Caco2 and HepG2 cells by analysis of hepatic transcriptome (RNAseq), western blot and ELISA. In vivo studies were conducted in C57BL/6J mice overexpressing mimic miR-100-5p.

**Results:** Postprandial variation in circ-miR-100-5p levels inversely correlate with PP-TG response. Cir-miR-100-5p was preferentially associated with TGRL particles of intestinal origin in subjects exhibited a low PP TG response. Differential analysis of transcriptome from HepG2 cells transfected by either mimic miR-100-5p or scrambled mimic miR as control allowed us to identify *PCSK9* as a down-regulated gene. Overexpression of miR-100-5p in HepG2 cells significantly decreased *PCSK9* mRNA levels by 52% (p<0.0001), cellular protein content by 28 % (p<0.0001) as well as PCSK9 secretion by 39% (p<0.0001). *In vivo* systemic delivery of mimic miR-100-5p induced a two-fold reduction (p<0.0001) on PP-TG in mice, such effect being abolished by blocking the circulating form of PCSK9 with alirocumab. Finally, we revealed a significant inverse relationship between circulating miR-100-5p expression levels and both PCSK9 levels and the magnitude of postprandial hypertriglyceridemia.

**Conclusion:** Taken together, our observations reveal that miR-100-5p regulates postprandial hypertriglyceridemia by targeting PCSK9, thus enhancing hepatic triglyceride-rich lipoproteins (TGRL) uptake. Our findings allow us to propose circ-miR-100-5p as a potential biomarker for early identification of subjects at high cardiovascular risk, prior to appearance of classical clinical features of metabolic disorders.

Postprandial clinical study, HDL-PP (NCT03109067)

**Lay summary:** This study examined whether miRs may control postprandial triglyceride response

*Key findings:* Our data reveal that miR-100-5p regulates postprandial hypertriglyceridemia by targeting PCSK9 Our observations allow us to propose miR-100-5p as a potential biomarker for early identification of subjects at high cardiovascular risk

## INTRODUCTION

Several epidemiological studies support that elevated non-fasting triglycerides (TG) concentrations are an independent risk factor for development of cardiovascular diseases (CVD)^1,2^. Based on these studies, a high postprandial triglyceride (PP-TG) response (non-fasting TG >2 mmol/l) has been associated with increased risk of CVD^3,4^. Since an elevated PP-TG response can manifest before clinical signs of metabolic disorders^5^, it may serve as a sensitive early marker for cardiovascular risk in apparently healthy individuals^6^. The link between postprandial TG metabolism and CVD is primarily attributed to the overproduction of triglyceride-rich lipoproteins (TGRL) and impaired clearance of their remnants^6^. Intravascular lipoprotein lipase (LPL)-mediated lipolysis of TGRL generates remnants particles which can contribute to CVD by generating pro-inflammatory, pro-coagulant and pro-oxidant products^7^. In addition, these remnants can penetrate the vascular intima, facilitating foam cell formation and accelerating atherosclerosis^8^. While some mechanisms underlying postprandial hypertriglyceridemia have been identified^9^, key factors driving exaggerated PP-TG elevations remain incompletely understood.

Recent evidence suggests that epigenetic mechanisms, particularly the regulation of gene expression by microRNAs (miRs)^10^, play a significant role in interindividual variability in PP-TG response^11^. miRs are small, non-coding RNAs, which can regulate gene expression at a posttranscriptional level, after binding to a complementary sequence of messenger RNA (mRNA) in the 3’ untranslated region, miRs either repress translation or cleave mRNA^12^. Mature miRs can be released from cells and be found in circulation associated with exosomes, apoptotic bodies, lipoproteins or bound to RNA binding proteins allowing for intercellular communications^13^. Notably, a large number of miRs have been identified as regulators of key genes involved in lipid and lipoprotein metabolism, making them interesting targets to study in the postprandial context^14–16^. Despite growing interest in miRs as regulators of lipid homeostasis, only few studies have explored their role in postprandial metabolism. One study demonstrated that postprandial TGRL isolated from human subjects after a moderately high-fat meal modulated miR-126 expression in human aortic endothelial cells^17^. Additional research using a Rainbow trout model observed changes in hepatic miR levels over the postprandial time course^18^ while mouse studies have shown that circulating miRs (circ-miRs) can be modulated by dietary fat intake^19,20^. In humans, analysis of peripheral blood mononuclear cells (PBMCs) following a high-fat meal revealed specific postprandial alterations in miR expression^21^, including modulation of circ-miR-92a levels^22^. Furthermore, distinct postprandial variations in circ-miRs have been observed between insulin-sensitive and insulin-resistant women, suggesting a potential role for miRs in metabolic inflexibility^23^. To date the contribution of miRs to the regulation of postprandial hypertriglyceridemia remains largely unexplored. Further investigations are needed to elucidate their role in lipid metabolism and their potential as biomarkers or therapeutic targets for cardiovascular disease prevention.

## METHODS

### Human fasting and postprandial plasma samples

Plasma samples were obtained from the previously described HDL-PP cohort^5^. The clinical HDL-PP study (NCT03109067) was carried out at the Clinical Investigation Center of the Pitié-Salpêtriere hospital (CIC-1901 Paris-Est). The institutional medical ethics committee approved the aforementioned clinical study and written informed consent was obtained from each participant. Clinical and biochemical characteristics of subjects are presented in **Table 1**. Each subjects underwent a postprandial exploration following consumption of a standardized solid test meal (freshly prepared commercially available foods: instant mashed potatoes (100g) mixed with 48g oil (2/3 sunflower oil and 1/3 rapeseed oil), beef steak (100g), cheese (28g), white bread (40g) and apple (120 g)) represented a total of 1200kcal and consisted of 14% protein (44g), 38% carbohydrates (119g) and 48% fat (66g)^24^. The healthy non-dyslipidemic population of male subjects was stratified according the degree of PP-TG response into two subgroups of subjects as follows: individuals with a low PP-TG response (iAUC-TG below median) and subjects with a high PP-TG response (iAUC-TG above median). None of the subjects had diabetes, liver disease, kidney disease or thyroid disease, nor were being treated with lipid-lowering drugs or medication affecting glucose homeostasis and all were non-smokers.

**Table 1.** Clinical and biological characteristic of the healthy non-dyslipidemic subjects stratified according postprandial TG response.

| Variables | All subjects<br>(n=32) | Subgroup with low PP-TG<br>response<br>(iAUC- TG below median) | Subgroup with a high PP-TG response<br>(iAUC- TG above median) |
| --- | --- | --- | --- |
| Age, year | 28.4±9.6 | 27.2±10.5 | 29.7±8.7 |
| BMI, kg/m<sup>2</sup> | 22.9±2.3 | 22.2±2.3 | 23.6±2.2 |
| Waist circumference, cm | 82.5±7.1 | 81.8±6.3 | 83.3±7.9 |
| SBP, mmHg | 118.5±7.9 | 116.6±8.9 | 120.5±6.4 |
| DBP, mmHg | 75.8±8.2 | 74.1±9.3 | 77.6±6.9 |
| FBG, mmol/l | 4.68±0.41 | 4.63±0.46 | 4.73±0.36 |
| HbA1c, % | 5.30 (5.10-5.40) | 5.20 (5.05-5.40) | 5.30 (5.20-5.50) |
| Insulin, mU/l | 1.75 (1.00-3.92) | 1.37 (1.00-4.38) | 2.05 (1.00-3.04) |
| HOMA-IR | 0.41 (0.20-0.83) | 0.32 (0.20- 0.94) | 0.43 (0.21-0.64) |
| TC, g/l | 1.69±0.34 | 1.57±0.31 | 1.82±0.32* |
| LDL-C, g/l | 1.06±0.29 | 0.94±0.25 | 1.17±0.28* |
| HDL-C, g/l | 0.50±0.13 | 0.52±0.12 | 0.48±0.14 |
| RLP-C, g/l | 11.6 (8.65-16.9) | 8.88 (7.67-13.1) | 15.07 (10.8-18.8)** |
| TG Before meal, g/l | 0.81 (0.59-1.18) | 0.59 (0.53-0.67) | 1.21 (1.0-1.35)** |
| TG 2 hours after meal, g/l | 1.45 (1.15-2.27) | 1.15 (0.96-1.39) | 2.44 (1.87-2.77)** |
| TG 4 hours after meal, g/l | 1.41 (1.14-2.27) | 1.14 (0.88-1.29) | 2.34 (1.86-2.99)** |
| TG 6 hours after meal, g/l | 1.17 (0.68-1.58) | 0.57 (0.69-0.94) | 1.57 (1.38 -1.95)** |
| AUC<sub>0-6h</sub>- TG, g/l.h | 7.97 (5.78-11.85) | 5.78 (5.21-6.42) | 12.08 (10.47-13.53)** |
| iAUC<sub>0-6h</sub> -TG, g/l.h | 3.04 (2.08- 5.09) | 2.08 (1.75-2.37) | 5.22 (4.08-6.12)** |

Values are mean±SD or median (interquartile range). BMI: Body Mass Index; SBP: Systolic blood pressure; DBP: Diastolic blood pressure; FBG: Fasting blood glucose; HOMA-IR: homeostasis model assessment of insulin resistance index; TC: Total cholesterol, RLP: Remnant lipoprotein; TG: Triglycerides; AUC: area under the curve; iAUC: incremental area under the curve.
\*p<0.05 and \*\*p<0.0001 versus subjects exhibiting a low PP-TG response

### Biochemical analyses

Biochemical measurements were performed with a calibrated autoanalyzer Konelab20 (Thermo Fisher Scientific, Courtaboeuf, France) using routine automated enzymatic methods. Triglycerides, total cholesterol, direct LDL-cholesterol, direct HDL-cholesterol and glucose were measured using Thermo Fisher Scientific commercial kits. Free cholesterol and phospholipids were measured by using reagents from Diasys (Holzheim, Germany). Insulin levels were determined using a human ELISA kit from Merck Millipore (Merck KGaA, Darmstadt, Germany). Insulin-resistance was evaluated using the homeostasis model assessment of insulin resistance index (HOMA-IR) using the following formula [fasting glucose (mmol/l) x fasting insulin (mUI/l)/22.5]. Bicinchoninic acid assay reagent from Pierce was utilized for total protein quantification (Thermo Fisher Scientific, Courtaboeuf, France). Human and murine PCSK9 measurements were performed using specific commercial ELISA Kits (R&D Sytems, Minneapolis, MN, USA) according to manufacturer’s instructions.

### Circulating microRNAs isolation and quantification

Circ-miRs were isolated from plasma or isolated lipoproteins with the miRNeasy Serum/Plasma kit (Qiagen, Hilden, Germany) according to the manufacturer’s guidelines. MicroRNAs from Caco2 and HepG2 cells or mice tissues were isolated with the miRNeasy Mini kit (Qiagen, Hilden, Germany) according to the manufacturer’s guidelines. Reverse transcription was performed using Human MegaPlex RT Primers Human Pool A v2.1 and Pool B v3.0 (Life technologies, Pleasanton, USA). Preamplification was performed using Human PreAmp Primers, Human Pool A v2.1 and Pool B v3.0 with TaqMan PreAmp Master Mix (Life technologies, Pleasanton, USA). For the discovery phase, a high-throughput array analysis was performed using TaqMan Array Human MicroRNA A+B Cards Set v3 and analyzed using the 7900HT Fast Real Time PCR System (Life technologies, Pleasanton, USA) to evaluate the expression profile of 754 human mature microRNAs. In further evaluations, identified circ-miRs levels were determined by quantitative stem-loop PCR technology with single TaqMan MicroRNA Assays (Life technologies, Pleasanton, USA) using a LightCycler 480 (Roche Diagnostics GmbH, Mannheim Germany). To calculate miR levels, Ct values were normalized to U6 small nuclear RNA endogenous control Ct values. Values are represented as relative quantitative value to U6 (RQV = 2^CtU6-CtmiR^), as fold change by applying the 2^-ΔΔCt^ method or as net Area Under the Curve for the Relative Quantitative Value to U6 (netAUC_0 to 6h_-RQV).

### Isolation and characterization of lipoproteins

Triglyceride-Rich Lipoprotein (TGRL) were isolated by centrifugation at 20,000 rpm for 45 min at 15°C using a SW41 Ti rotor in a Beckman XL70 ultracentrifuge (Beckman Coulter). Low-density lipoprotein (LDL) and High-density lipoprotein (HDL) were isolated from TGRL-free plasma by a two-step ultracentrifugation procedure at 100 000 rpm using a Beckman TLA100.3 rotor in a Beckman Optima Max TL ultracentrifuge at 15°C. Briefly, the density of the TGRL free plasma was adjusted to d>1.063 g/ml fraction and centrifuged 3h30 to collect LDL fraction. Then, HDL fraction (d 1.063- 1.21 g/ml) were obtained following a second ultracentrifugation for 5h30.

### Differentiation and transfection of human Caco2 cells

Caco-2 cells, human colorectal epithelial adenocarcinoma (ECACC), were grown in Dulbecco’s Modified Eagle’s Medium - high glucose (DMEM; Sigma-Aldrich) containing 20% fetal bovine serum (FBS; Sigma-Aldrich), 1% L-glutamine solution (Sigma Aldrich), 1% Penicillin-Streptomycin (Sigma-Aldrich), 1% MEM Non-essential Amino Acid solution, and 1% Sodium pyruvate solution at 37°C in 5% CO2. The cells were cultured and used when they reached 80% confluence. The cells were seeded on Transwells® (24 mm diameter, 3 μm pore size, polyester (PET) membrane, Corning Costar Corp, Cambridge, MA), and the medium was changed every two days until differentiation (21 days) as previously described^25^. Cells were transfected using Lipofectamine RNAiMAX (Invitrogen) with either hsa-miR-100-5p mirVana miRNA mimic (mimic miR 100-5p; Invitrogen) or negative control #1 mirVana miRNA mimic (mimic Scrambled; Invitrogen) for 24 hours, according to the manufacturer’s instructions. Then, cells were incubated with FBS-free culture medium enriched with lipid cocktail (TLO: sodium taurocholate [1 mmol/L], lecithin [0.3 mmol/L], and oleic acid (OA) [1.6 mmol/L]) for 4 hours in the apical chamber (to induce the secretion of triglyceride-rich lipoproteins) and with FBS-depleted culture medium in the basal chamber as previously described^25,26^. To isolate the particles, the basal medium containing the lipoproteins was subjected to ultracentrifugation as described above.

### Transfection of human HepG2 cells

Human hepatoma HepG2 cells (ATCC, Manassas, VA, USA) were grown in Dulbecco′s Modified Eagle′s Medium - high glucose (DMEM; Sigma-Aldrich, United Kingdom) containing 10% fetal bovine serum (FBS; Sigma-Aldrich), 1% L-Glutamine solution (Sigma-Aldrich) and 1% Penicillin-Streptomycin (Sigma-Aldrich) at 37°C in 5% CO2. Cells were plated and used when 80 % confluent. Cells were transfected by using Lipofectamine RNAiMAX (Invitrogen) with either hsa-miR-100-5p mirVana miRNA mimic (mimic miR-100- 5p; Invitrogen) or negative control #1 mirVana miRNA mimic (mimic Scrambled; Invitrogen) for 24h according manufacturer’s instructions.

### RNA extraction, RNA sequencing, reverse transcription, and quantitative PCR

Total RNA was extracted from HepG2 cells with the miRNeasy Mini kit (Qiagen, Hilden, Germany) according to the manufacturer’s protocol. A total of 1000 ng of RNAs per sample was used to prepare the mRNA library, followed by cluster amplification and sequencing. The concentration and size distribution of all samples were determined by 4200 TapeStation (Agilent Technologies). Polyadenylated RNA transcripts were selected for library preparation using oligo (dT) primers attached magnetic beads, then RNA was fragmented before random priming. First and second strand cDNA were synthesized before end-repair, followed by the addition of a single ’A’ nucleotide to the 3’ ends of the blunt fragments. Index adapters were ligated to the 5’ and 3’ ends, preparing the double-stranded cDNA for hybridization onto a flow cell. Afterwards, cluster generation was achieved as each fragment was amplified into clonal clusters via bridge amplification. Next-generation Sequencing was then performed in a paired-ended mode, 2 x 51 bp, on a NovaSeq™ 6000 Sequencing System (Illumina) with the NovaSeq 6000 SP Reagent Kit (100 cycles) following the manufacturer’s protocols. Read qualities were assessed using Trimmomatic and FastQC. Global mapping was established with STAR to the human genome (GRCh37/hg19). The reads count per gene was achieved using RSEM software. Normalization was done on the size of libraries. This work benefited from equipment and services from the iGenSeq core facility at ICM (Institut du Cerveau et de la Moelle Epinière, Paris, France). Bioinformatic analyses were conducted by the Data and Analysis Core platform at ICM, providing a graphical Shiny/R application for exploring results. Conducting supervised analysis, differentially expressed genes were considered statistically significant with an adjusted p-value < 0.05 with Benjamini & Hochberg False Discovery Rate method. Gene Set Enrichment Analysis (GSEA) was performed using the Hallmark Gene Set Collection.

Relative quantification of target mRNA was performed using specific primers on LightCycler 480 (Roche Diagnostics GmbH, Mannheim Germany) and was normalized to the mean expression of three different housekeeping genes for human or mice: Hypoxanthine-guanine phosphoribosyltransferase (HPRT1), NonPOU domain-containing octamer-binding protein (NONO) and Tubulin alpha-1A chain (TUBA1A) for samples obtained from HepG2 cells; Hypoxanthine-guanine phosphoribosyltransferase (Hprt), NonPOU domain-containing octamer-binding protein (NONO) and beta-glucuronidase (β-gus) for mouse samples. Data were expressed as a relative change in mRNA expression or a fold change compared to control values.

### Protein extraction and Western blotting

Caco2 or HepG2 transfected cells were lysed on ice using M-PER Mammalian Protein Extraction Reagent (Thermo Fisher Scientific) supplemented with cOmplete^TM^, Mini Protease Inhibitor Cocktail (Roche Diagnostics GmbH). Lysates were then centrifuged at 14000g for 10 min at 4°C to remove cell debris. Protein concentrations of each sample were measured using Pierce BCA Protein Assay Kit (Thermo Fisher Scientific), following the manufacturer’s guidelines. Samples were boiled at 95°C for 5 min and proteins (20 µg/lane) were separated under reducing conditions on a NuPAGE 4 to 12%, Bis-Tris, 1.0–1.5 mm, Mini Protein Gels (Invitrogen) in an electrophoresis running buffer with NuPAGE MES SDS Running Buffer 1X (Invitrogen). Proteins were transferred onto nitrocellulose membranes (0.45 µm, Amersham) and blocked in Blocker Casein in PBS (Thermo Fisher Scientific) for 1 h at room temperature. Membranes were then incubated overnight with the specific primary antibody (Human PCSK9 antibody, AF3888, Bio-techne; or Anti-β-Actin antibody, A5441, Sigma-Aldrich) in Blocker Casein in PBS at 4°C overnight. After washing and incubation, membranes were labeled with HRP-conjugated secondary antibody (Sheep IgG Horseradish Peroxidase conjugated Antibody, HAF016, Bio-techne) or a fluorescent-conjugated secondary antibody (IRDye® 800CW Goat-anti-Mouse Antibody, 926-32210, LI-COR) for 1 h at room temperature in Blocker Casein in PBS. Membranes were then washed, developed and captured digitally using a LI-COR Odyssey Infrared Imaging System 9120 (for the fluorescent secondary antibodies) or revealed by enhanced chemiluminescence by incubating the membrane with Pierce ECL Western Blotting Substrate (Thermo Fisher Scientific) coupled with an exposure on the ImageQuant LAS 4000 (GE Healthcare) for the HRP-conjugated secondary antibodies. Bands intensities were assessed by ImageJ and expressed relative to β-actin.

### Animal experiments

#### Mouse strain, animal housing and diets

Nine-weeks old C57BL/6J mice obtained from Janvier Labs (Le Genest St Isle, France) were randomly assigned to experimental groups and fed a regular chow diet (CD; A04-10, Safe lab, Rosenberg, Germany). Mice were maintained on a 12 h light and dark cycle at 22°C with ad libitum access to water and diet in specific pathogen-free conditions in the animal facility at Sorbonne University. All animal procedures were performed in accordance with the guidelines of the Charles Darwin Ethics Committee from Sorbonne University on animal experimentation and with the French Ministry of Agriculture (APAFIS # 20376-2019042514251937). Moreover, all animal procedures involving mice were carried out according to the Guide for the Care and Use of Laboratory Animals published by the European Commission Directive 2010/63/EU revising directive 86/609/EEC.

#### miRNA injections

miRIDIAN stabilized mimic of hsa-miR-100-5p (mimic miR-100-5p; MIMAT0000098, DharmaconTM, Horizon discovery) were used for in vivo delivery in mice. Mimic was delivered weekly via retro-orbital injections, with a dose of 1.5 nmol in association with a transfecting reagent (in vivo-jetPEI®, Polyplus transfection). For the control group we used a miRIDIAN stabilized mimic Scrambled (mimic scrambled; cel-miR-67-3p, MIMAT0000039, DharmaconTM, Horizon discovery).

#### Evaluation of postprandial hypertriglyceridemia in mice

Oral Lipid Tolerance Tests (OLTT) were performed as follows: mice were fasted overnight for 12 hours before OLTT. Fasted mice were orally administrated 200µl olive oil. Blood was collected from the tail vein before and at 1, 2, 3, 4, 6 and 8 hours after gavage. Plasma TG levels were measured and incremental area under the curve (iAUC) value of plasma TG levels was calculated using the trapezoidal rule.

When indicated OLTT were equally performed 3 days after subcutaneous injection of monoclonal antibody against PCSK9 as previously described^27^. Alirocumab (Praluent, Sanofi-Aventis France) was diluted in saline solution (NaCl 0,9%) and administrated subcutaneously at a dose of 10 mg/kg of body weight.

### Data analysis and statistics

Variables were tested for normal distribution using the Kolmogorov–Smirnov test. Circulating levels of miRs did not follow a Gaussian frequency distribution. The Friedman test was used to assess changes over the postprandial time course. The Dunn’s test was applied for multiple comparisons. Postprandial triglyceride variations were quantified by calculating the incremental AUC (iAUC). Differences between two groups were analyzed using an unpaired non-parametric Mann-Whitney test. Statistical analyses were performed using either GraphPad Prism 8. Results were considered statistically significant at p < 0.05.

## RESULTS

### Identification of circulating miRs associated with postprandial triglyceride response

In order to identify circ-miRs that vary during the postprandial phase, we compared circ-miRs levels of 5 healthy non-dyslipidemic subjects’ representative of our study population, at fasting (T0) and 2 hours after the meal intake (T2), by high-throughput array analysis. Among detected miRs, we identified 15 circ-miRs with increased levels during the postprandial state, with a fold change greater than 1.6 between T2 and T0 (**Supplemental Table S1**). We next assessed levels of identified circ-miRs throughout the postprandial phase in 32 healthy non-dyslipidemic subjects. A significant increase in plasma circ-miRs levels was observed specifically in subjects exhibited a low postprandial TG response (iAUC-TG below the median) **(Table 2)**. Correlation analyses reveal that for most of identified miRs, postprandial variation in circ-miR levels were inversely associated with the degree of postprandial triglyceride response **(Table 2),** with the strongest association observed for circ-miR-100-5p (r=0.4137; p<0.0001) **(Figure 1A). Figure 1B** shows differential postprandial variation in circ-miR100-5p levels among non-dyslipidemic healthy individuals stratified according the degree of postprandial TG response. Quantification of postprandial circ-miR variation by calculation of the net incremental Area Under the Curve for the Relative Quantitative Value to U6 (nAUC_0 to 6h_-RQV) reveals that individual exhibited a high postprandial TG response were characterized by significant reduced postprandial circ-miR100-5p levels as compared to those exhibited a low postprandial TG response **(Figure 1C).** Notably, similar fasting levels of circ-miR were observed in subjects exhibited either a low or a high postprandial TG response. **(Supplemental Figure S1).**

**Figure 1.**
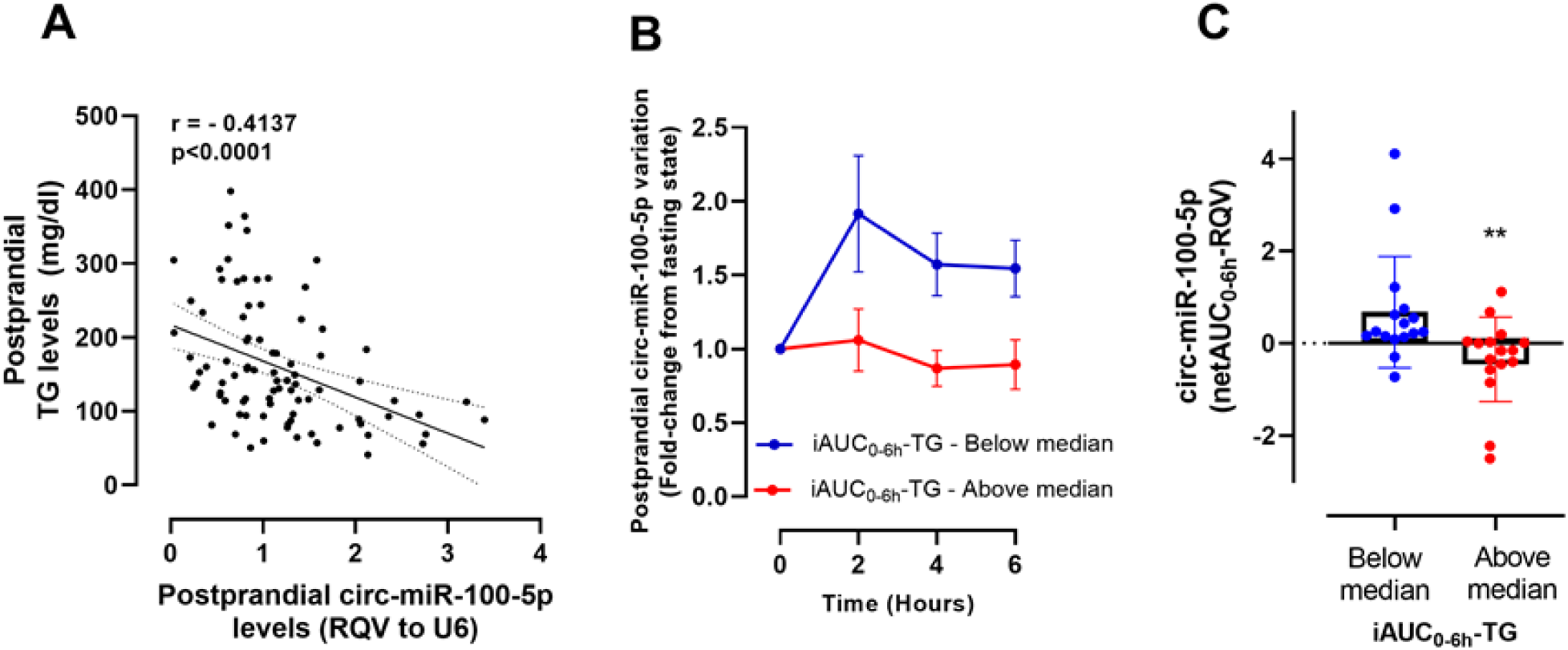
Differential circ-miR-100-5p variation according the degree of postprandial TG response. (**A)** Inverse relationship between postprandial variation in circ-miR-100-5p and triglycerides levels in non-dyslipdemic subjects (n=32). Postprandial circ-miR100-5p variation expressed as **(B)** fold change from fasting state and as**(C)** net incremental Area Under the Curve for the Relative Quantitative Value to U6 (nAUC_0→6h_-RQV) in non-dyslipdemic subjects exhibited a low PP-TG response (iAUC-TG below median, n=16) or a high postprandial TG response (iAUC-TG above median, n=16). **p<0.005.

**Table 2.** Fold changes in postprandial circulating miRs levels as a function of postprandial TG response in healthy non-dyslipidemic population.

| miRNA | Hours | Subgroup with low<br>PP-TG response<br>(iAUC- TG below<br>median) | Subgroup with a high<br>PP-TG response<br>(iAUC- TG above<br>median) | Relationship between<br>postprandial changes in<br>circ-miR and TG levels |  |
| --- | --- | --- | --- | --- | --- |
|  |  |  |  | r | p |
| hsa-miR-1-3p | 2 | **3.18±1.17\*** | 1.82±0.36 | -0.1414 | 0.1740 |
|  | 4 | **2.77±0.71\*** | 1.50±0.36 |  |  |
|  | 6 | 2.40±0.59 | 1.44±0.24 |  |  |
| hsa-miR-100-5p | 2 | **1.91±0.39\*** | 1.06±0.21 | -0.4135 | <0.0001 |
|  | 4 | **1.57±0.21\*** | **0.87±0.12\*** |  |  |
|  | 6 | **1.54±0.19\*** | **0.89±0.17\*** |  |  |
| hsa-miR-107 | 2 | **2.05±0.49\*** | 1.14±0.22 | -0.3807 | 0.0002 |
|  | 4 | 1.73±0.31 | 0.99±0.19 |  |  |
|  | 6 | 1.82±0.27 | 1.02±0.22 |  |  |
| hsa-miR-139-5p | 2 | 1.30±0.16 | 1.04±0.16 | -0.1574 | 0.1298 |
|  | 4 | 1.43±0.17 | 0.81±0.10 |  |  |
|  | 6 | 1.37±0.23 | 0.89±0.10 |  |  |
| hsa-miR-140-3p | 2 | **1.73±0.14\*** | 1.34±0.24 | -0.3316 | 0.0010 |
|  | 4 | **1.89±0.20\*** | 1.19±0.19 |  |  |
|  | 6 | 1.64±0.22 | 1.15±0.15 |  |  |
| hsa-miR-154-3p | 2 | **2.71±0.62\*** | 1.15±0.34 | -0.3740 | 0.0005 |
|  | 4 | 2.10±0.50 | 0.94±0.39 |  |  |
|  | 6 | 1.59±0.26 | 1.19±0.44 |  |  |
| hsa-miR-16-5p | 2 | **2.18±0.54\*** | 1.55±0.34 | -0.3235 | 0.0012 |
|  | 4 | **1.82±0.14\*** | 1.12±0.14 |  |  |
|  | 6 | 1.68±0.21 | 1.26±0.25 |  |  |
| hsa-miR-16-1-3p | 2 | 1.77±0.25 | 1.16±0.20 | -0.3239 | 0.0013 |
|  | 4 | **1.98±0.33\*** | 0.88±0.12 |  |  |
|  | 6 | 1.37±0.23 | 0.97±0.15 |  |  |
| hsa-miR-26b-5p | 2 | 1.56±0.21 | 1.56±0.28 | -0.2960 | 0.0085 |
|  | 4 | **1.78±0.23\*** | 1.60±0.48 |  |  |
|  | 6 | 1.84±0.31 | 1.04±0.182 |  |  |
| hsa-miR-29a-5p | 2 | **2.34±0.61\*** | 1.33±0.39 | -0.3186 | 0.0028 |
|  | 4 | 1.93±0.27 | 0.93±0.19 |  |  |
|  | 6 | 1.22±0.19 | 1.11±0.31 |  |  |
| hsa-miR-29b-2-5p | 2 | **2.77±0.54\*** | 1.29±0.20 | -0.2689 | 0.0146 |
|  | 4 | 2.24±0.43 | 1.23±0.26 |  |  |
|  | 6 | 2.34±0.50 | 1.11±0.23 |  |  |
| hsa-miR-431-5p | 2 | 1.84±0.56 | 1.97±0.46 | -0.1217 | 0.2585 |
|  | 4 | 1.61±0.21 | 2.13±0.68 |  |  |
|  | 6 | 1.35±0.17 | 1.19±0.20 |  |  |
| hsa-miR-744-3p | 2 | **1.80±0.31\*** | 1.18±0.20 | -0.3323 | 0.0020 |
|  | 4 | **2.30±0.55\*** | 1.09±0.26 |  |  |
|  | 6 | 1.63±0.24 | 1.10±0.2 |  |  |
| hsa-miR-9-5p | 2 | **1.92±0.30\*\*** | 1.12±0.29 | -0.2871 | 0.0070 |
|  | 4 | 1.37±0.18 | 0.80±0.19 |  |  |
|  | 6 | 1.39±0.30 | **0.64±0.12\*** |  |  |
| hsa-miR-941 | 2 | 1.86±0.33 | 1.09±0.15 | -0.2946 | 0.0069 |
|  | 4 | **2.25±0.41\*** | 0.95±0.18 |  |  |
|  | 6 | 1.72±0.25 | 0.80±0.16 |  |  |

\*p<0.05 adjusted p value vs before test meal

### Characterization of TGRL-associated miR-100-5p according to the degree of postprandial TG response

Transient elevation of plasma TG levels following meal intake reflects the appearance of postprandial TGRL into the intravascular compartment. As expected, individuals exhibiting a high PP TG response displayed higher circulating levels of TGRL-TG as compared to subjects with a low PP TG response **(Figure 2A).** Moreover, TGRL particles isolated from individuals from the high PP TG response subgroup were significantly enriched in TG compared to their counterpart isolated from the subjects with a low PP TG response **(Figure 2B).** As postprandial elevation in circ-miR-100-5p levels observed in healthy non-dyslipidemic subjects exhibited a low PP-TG response occurred concomitantly with those of TG levels, we thus evaluated whether circ-miR100-5p may be associated with circulating lipoproteins and more specifically with postprandial TGRL. Circ-miR-100-5p were detected on each of lipoprotein fractions, *i.e.* TGRL, LDL and HDL. To evaluate miR-content per lipoprotein particles, miR RQV to U6 was expressed to apolipoprotein B48 mass of TGRL particles. We observed an increased content in miR-100-5p in postprandial TGRL particles as compared those isolated from fasting samples (**Figure 2C**). More strikingly, we detected a specific higher content in miR-100-5p (4.5-Fold) in postprandial TGRL particles isolated from subjects exhibiting a low postprandial TG response as compared to their counterparts isolated from subjects exhibiting a high postprandial TG response.

**Figure 2.**
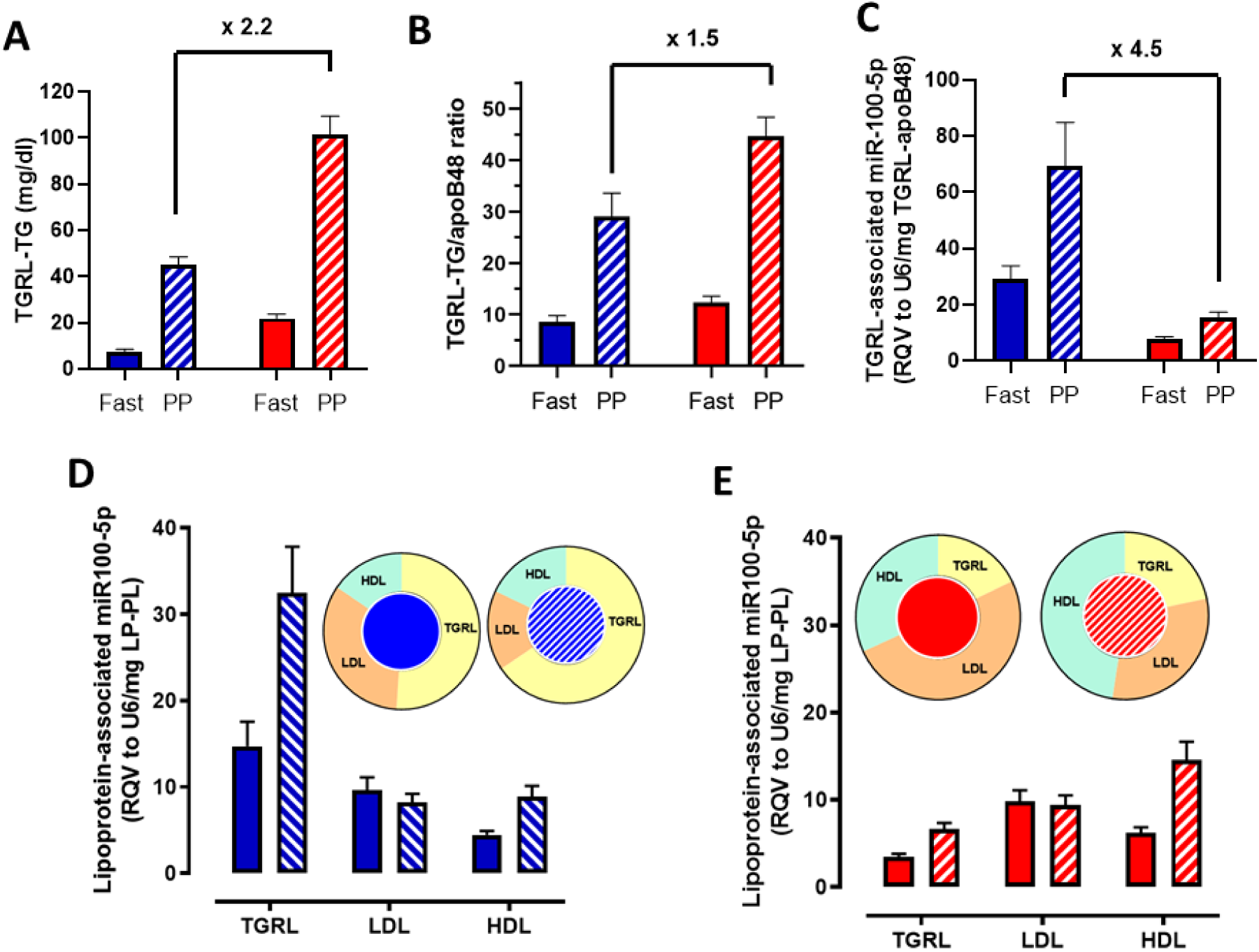
Differential lipoprotein-associated miR-100-5p levels according to the degree of postprandial TG response. (**A**) TGRL-TG levels, **(B)** TGRL-TG content and **(C**) TGRL-miR-100-5p content determined in fasting state (solid bars) and at the postprandial peak for each individual (hatched bars) exhibiting a low (iAUC below median, blue bars) or a high (iAUC-TG above median, red bars) postprandial TG response. Lipoprotein-associated miR-100-5p, expressed RQV to U6 per mg of phospholipid content, in TGRL, LDL and HDL in fasting (solid bars) and postprandial (hatched bars) states from subjects exhibiting **(D)** a low (blue bars) or **(E)** a high (red bars) postprandial TG response. Inserts in panels D and C show distribution of miR-100-5 p (expressed as percentage) among major lipoprotein classes in both fasting and postprandial states. Fast: fasting state; PP: postprandial state. Values are mean ± SEM.

In order to compare lipoprotein-associated miR levels between the different lipoprotein classes, we used the RQV to housekeeping gene U6 relative to the lipoprotein-phospholipid content, as phospholipid content represent the sole component with roughly an equivalent relative proportion between each class of lipoproteins^28^. Individuals exhibited a low postprandial TG response, circ-miR-100-5p was preferentially associated with TGRL particles, in both fasting and postprandial states **(Figure 2E),** whereas LDL and HDL particles represented preferential carriers of circ-miR-100-5p in subjects with a high postprandial TG response **(Figure 2D)**. These observations indicate that an elevated postprandial TG response is associated not only with overall reduced content of cir-miR-100-5p of TG enriched TGRL particles but equally with a differential distribution of miR-100-5p among lipoprotein particles.

To examine whether intestinal cells may contribute to formation and secretion of TGRL-associated miR-100-5p particles, differentiated Caco2 cells were transfected by either mimic miR-100-5p or scrambled mimic as control (**Figure 3A**), leading to an increase in miR100-5p level compared with control scrambled miR (**Figure 3B**). A similar amount of TGRL particles were isolated from basolateral media obtained from cells independently of miR-100-5p expression level (**Figure 3CD**). However, TGRL particles secreted from cells transfected by mimic-miR100-5p were characterized by a significantly reduced TG content (**Figure 3E**) and were enriched in miR100-5p (**Figure 3FG**) as compared to those isolated from cells transfected mimic-miR scrambled.

**Figure 3.**
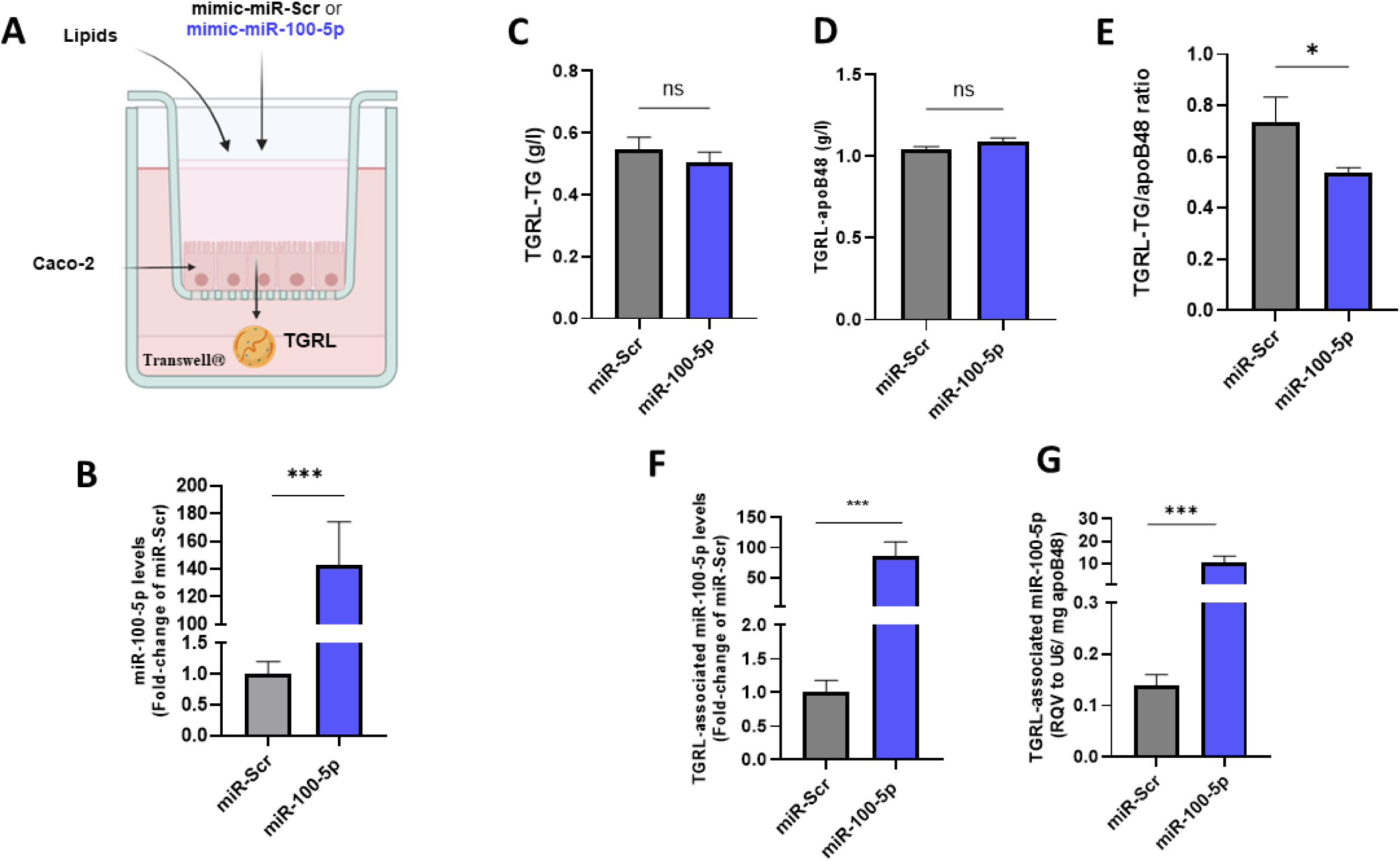
Impact of intestinal in vitro overexpression of miR-100-5p on TGRL secretion. **(A)** Experimental design for Caco2 cell differentiation, transfection and TGRL secretion. **(B)** miR-100-5p expression levels expressed as fold-change of miR-Scrambled determined following transfection of differentiated Caco2 cells with either miR-100-5p or a control mimic Scrambled (miR-Scr). Characterization of TGRL particles isolated from the basolateral media from cells transfected by mimic miR-100-5p or control mimic Scrambled (miR-Scr): **(C)** TGRL-TG levels, **(D)** TGRL-apoB48 levels, **(E)** TGRL-TG content expressed as TGRL-TG/apoB48 ratio, **(F)** TGRL-miR100-5p levels and **(G)** TGRL-miR-100-5p content, expressed as TGRL-miR100-5p/apoB48 ratio. Values are mean ± SEM. *p<0.05 and *** p< 0.0001.

Following LDL-mediated TG hydrolysis, chylomicron are progressively transforms into smaller remnant particles that are subsequently removed from the circulation by interaction with their specific hepatic receptors^29^. As lipoprotein-associated miRs can be deliver to target cells and regulate gene expression^13^, our observation suggest that individuals characterized by a low PP TG response exhibit an increased lipoprotein-associated miR-100-5p hepatic delivery as compared to subjects with a high PP TG response.

### Identification of PCSK9 as a target of miR-100-5p

To investigate the molecular mechanisms underlying the control of postprandial hypertriglyceridemia by miR-100-5p, by conducting an unbiased analysis of hepatic transcriptome (RNAseq) of HepG2 cells transfected by either mimic miR-100-5p or scrambled mimic as control **(Figure 4A).** Differential analysis identified eight down-regulated genes and four up-regulated genes in the group transfected with mimic miR100-5p compared to the control group **(Figure 4B).** Among those genes, we identified *PCSK9* as a downregulated gene of miR-100-5p. *In silico* analysis of miRNA target prediction databases^30^ revealed that PCSK9 is a potential target of miR100-5p (miRDB; http://mirdb.org). Additionally, using the web tool miRNA-centric network analytics miRNet (http://mirnet.ca), an interaction between miR-100-5p and *PCSK9* was also observed^31^. Furthermore, gene set enrichment analysis (GSEA) revealed that overexpression of miR-100-5p was associated with ten significantly regulated pathways, with the “cholesterol_homeostasis” as one of the most regulated pathways **(Figure 4C).** Then, we confirmed by qRT-PCR that overexpression of miR-100-5p in HepG2 cells significantly decreased h*PCSK9* mRNA levels by 52% (p<0.0001, **Figure 4D**). We equally observed a significant decrease in hPCSK9 protein content in miR-100-5p transfected cells (−28% p<0.0001; **Figure 4E**). Finally, overexpression of miR-100-5p in HepG2 cells significantly decreased hPCSK9 secretion by 39% (p<0.0001; **Figure 4F**), thus suggesting that miR-100-5p contributes to the regulation of circulating levels of PCSK9.

**Figure 4:**
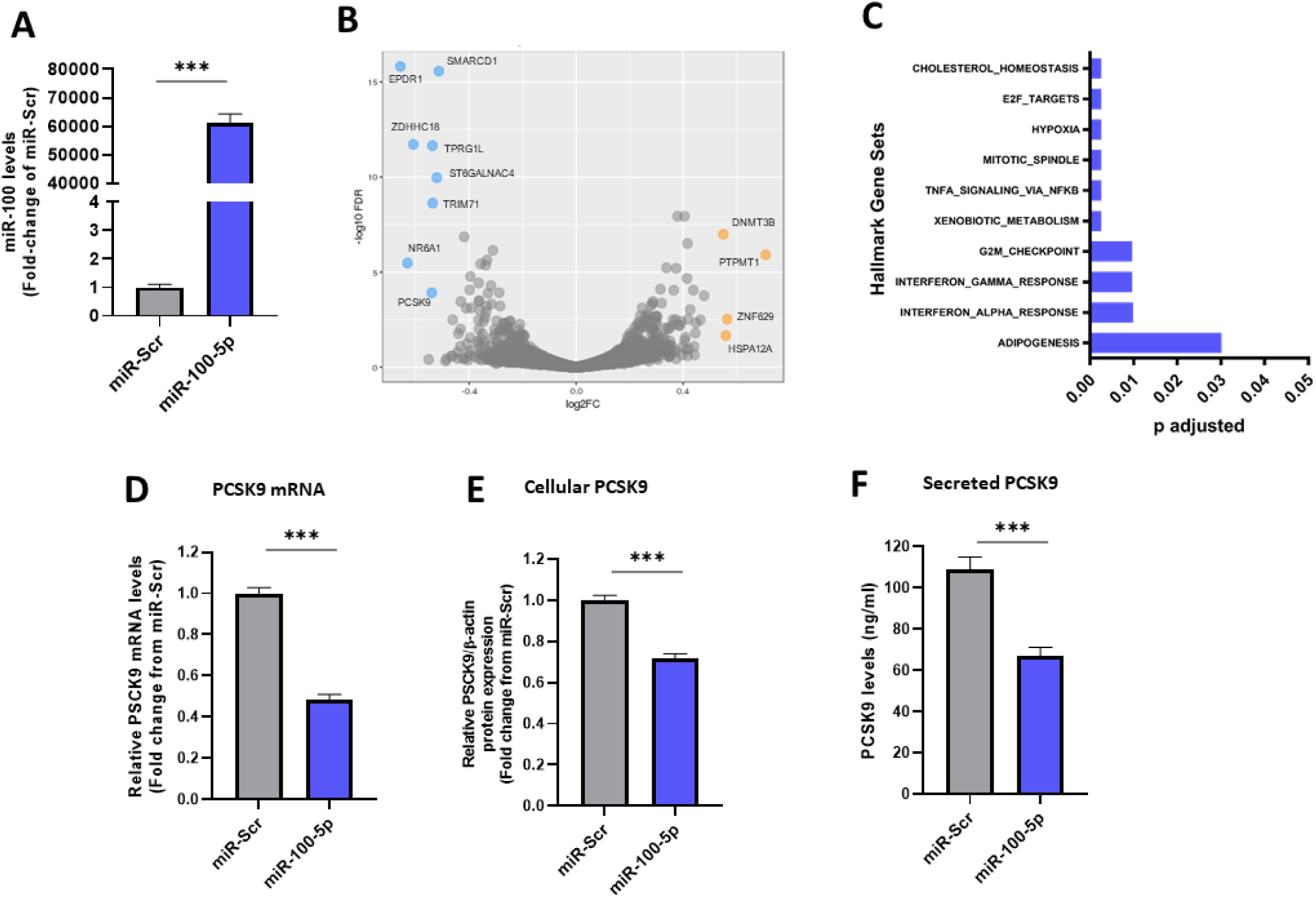
Impact of hepatic in vitro overexpression of miR-100-5p on PCSK9 expression and secretion. **(A)** miR-100-5p expression levels expressed as fold-change of miR-Scrambled determined following transfection of HepG2 cells with either miR-100-5p or a control mimic Scrambled (miR-Scr). (**B**) Volcano plot depicting the log2 fold change and corresponding - log10 adjusted p-value of all genes detected in HepG2 cells transfected with mimic miR-100-5p compared to control, with mimic miR-Scrambled as control. Differentially expressed genes (False Discovery Rate < 0.05) are shown in color (blue, down-regulated in miR-100-5p condition; orange, up-regulated in miR-100-5p condition). The log2 fold-change threshold for significant differences is set at 0.5 (FC > 1.4). Genes that did not have significant differences in expression are shown in grey. (**C**) Gene Set Enrichment Analysis using the Molecular Signatures Database Hallmark Gene Set Collection showing gene sets that are the most highly regulated. (**D**) mRNA levels of PCSK9, (**F**) cellular PCSK9 protein and (**G**) extracellular secreted PCSK9 levels determined following transfection of HepG2 cells with either miR-100-5p or a control mimic Scrambled. Values are mean ± SEM. *** p< 0.0001.

### Overexpression of miR-100-5p reduces postprandial hypertriglyceridemia in vivo

The experimental design for the evaluation of the impact of weekly injections of miR-100-5p on postprandial triglyceride response in C57BL/6 mice is depicted in a flow chart **(Figure 5A**). The systemic delivery of mimic miR-100-5p in mice led to hepatic overexpression of miR-100-5p (**Figure 5B**), and in a lesser extend in the intestine specifically in the jejunum (**Figure 5C**), in comparison to the control group injected with a control mimic-miR scrambled, as well as increased levels in plasma **(Figure 5D)**. After 4 weeks of treatment, we observed a two-fold reduction in postprandial TG response **(Figure 5EF**) in mice overexpressing mimic miR-100-5p as compared to those receiving weekly injection of mimic-miR scrambled. Notably, as shown in **Supplemental Figure S2**, mice overexpressing mimic-miR-100-5p exhibited a maximal reduction in postprandial TG response at 4 weeks that was maintained up to 8 weeks, whereas no significant change in postprandial TG response was observed in control mice throughout the 8-weekly injection of mimic-miR-scrambled. In addition, in good agreement with our in vitro analyses, we observed a significant reduction in circulating levels of mPCSK9 (21%; p=0.032) in mice overexpressing miR-100-5p (**Figure 5G)** together with a reduction of plasma cholesterol levels (**Figure 5H)** as a result of reduction in hepatic mPCSK9 gene expression **(Figure 5I).**

**Figure 5.**
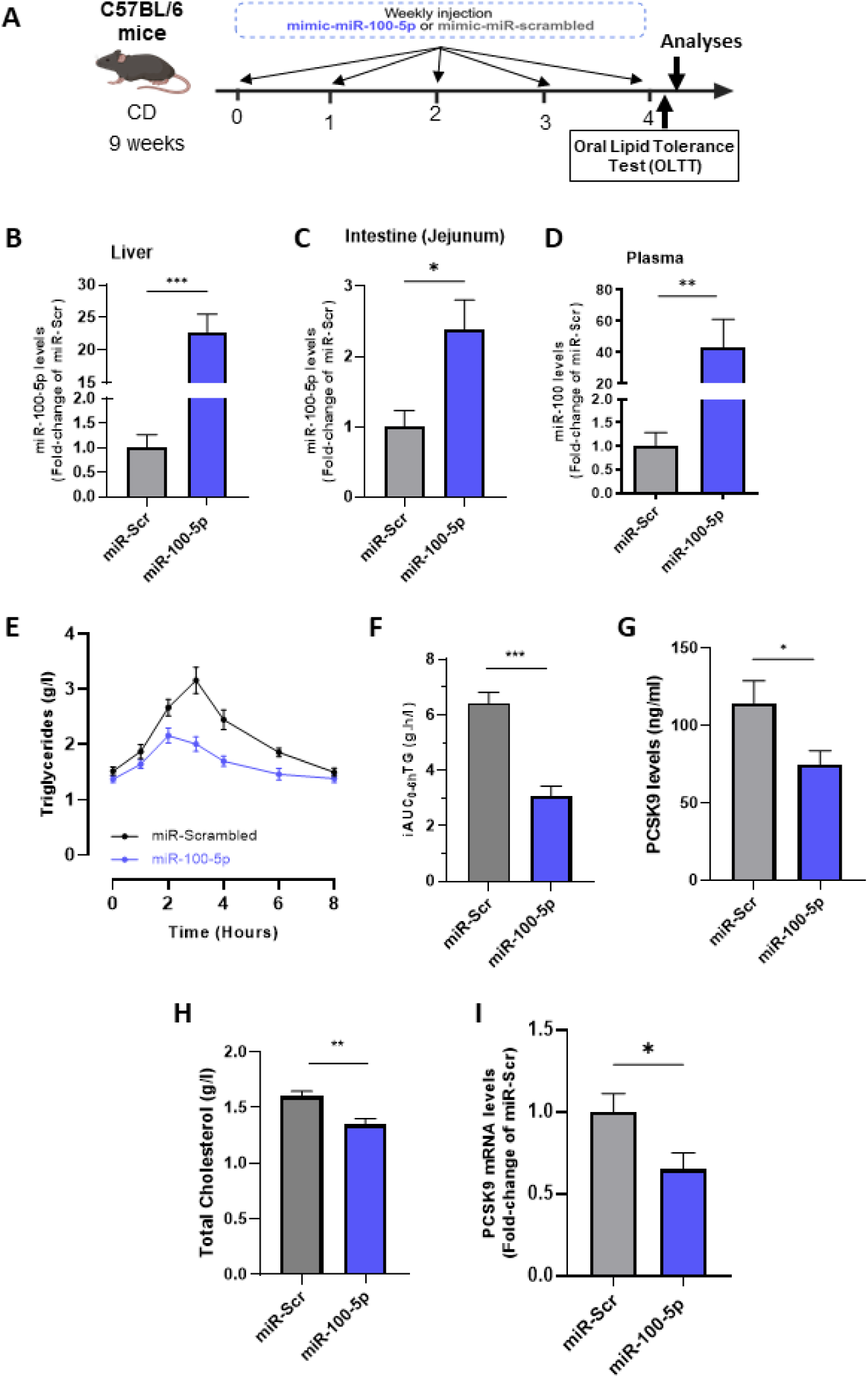
Impact of overexpression of miR-100-5p on postprandial hypertriglyceridemia in mice. **(A)** Experimental design, **(B)** Liver, **(C)** intestine and **(D)** plasma miR-100-5p levels in mimic-scrambled (n=16) or miR-100-5p (n=15) injected mice fed a chow diet. **(E)** Time course of triglycerides levels during OLTT determined after 4 weeks of weekly injection of mimic miR-Scrambled (black line) or miR-100-5p (blue line). **(F)** iAUC-TG_0→8h_ determined after 4 weeks of weekly injection in mimic-scrambled or miR-100-5p injected mice **(G).** Circulating PCSK9 and **(H)** plasma cholesterol levels determined in mice after 4 weeks of weekly injection of mimic miR-Scrambled or miR-100-5p. **(G)** hepatic mRNA levels of mPCSK9 after 4 weeks of weekly injection of mimic miR-Scrambled or miR-100-5p. Values are mean ± SEM. * p< 0.05; ** p<0.01 and ***<p<0.0001. iC: incremental area under the curve; TG: Triglycerides; OLTT: Oral lipid tolerance test.

To confirm that miR100-5p regulates postprandial TG response through its action on circulating levels of PCSK9, postprandial lipemia was determined three days after subcutaneous injection of vehicle or human PCSK9 monoclonal antibody (alirocumab, 10mg/kg) to capture circulating PCSK9 **(Figure 6A),** as previously described^27^. As expected, Alirocumab significantly reduced plasma cholesterol levels by approximately 15% compared with vehicle either in mice overexpressing miR-100-5p control mice injected with mimic-miR scrambled **(Figure 6B).** Alirocumab treatment significantly reduced postprandial TG response in control mimic-miR scrambled to a similar extend as that observed in mice overexpressing mimic miR-100-5p but was without any significant impact in those latter mice **(Figure 6BC).**

**Figure 6:**
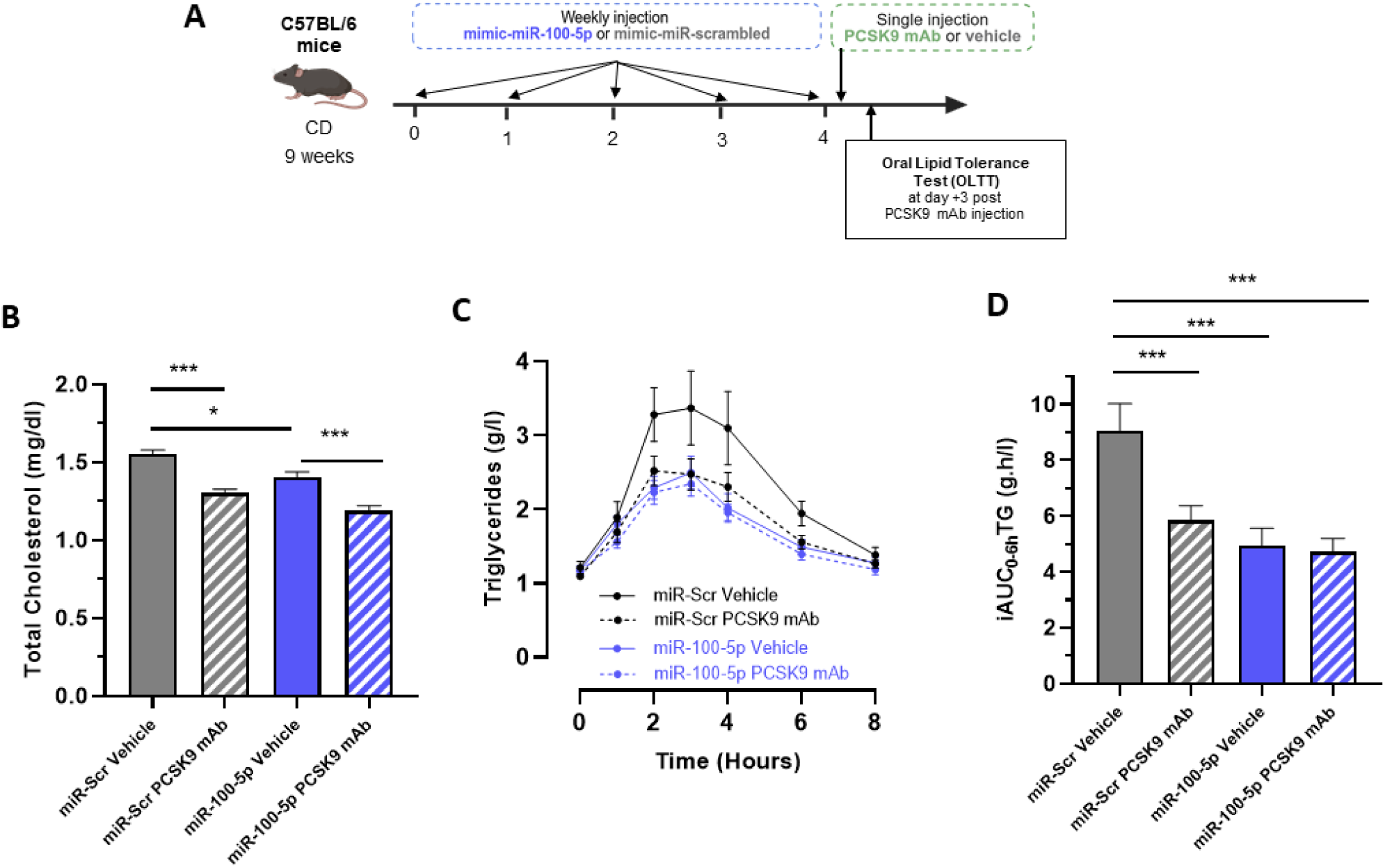
Overexpression of miR-100-5p reduces postprandial TG response through its action on circulating PCSK9. **(A)** Experimental design. Mice overexpressing mimic miR-100-5p received a subcutaneous administration of PCSK9 antibody (n=9, Alirocumab, 10 mg/kg) or a single subcutaneous injection of vehicle (n=8) and control mimic-miR scrambled mice received a subcutaneous administration of PCSK9 antibody (n=8, Alirocumab, 10 mg/kg) or a single subcutaneous injection of vehicle (n=7). Three days later, **(B)** plasma cholesterol levels were assessed after overnight fasting and **(C)** plasma triglycerides levels were determined during OLTT, time course of triglycerides levels during OLTT, **(D)** iAUC-TG_0→8h_. Values are mean ± SEM. *p<0.05 and ***<p<0.0001. iAUC: incremental area under the curve; TG: Triglycerides; OLTT: Oral lipid tolerance test.

### Relationship between circulating miR-100-5p expression and plasma levels of PCSK9

Consistent with our *in vitro* and *in vivo* observations, non-dyslipidemic individuals who exhibited a low postprandial TG response (iAUC-TG below median) were characterized by reduced plasma hPCSK9 levels compared to non-dyslipidemic subjects who exhibited a high postprandial TG response (iAUC-TG above median) **(Figure 7A).** In addition, we observed a strong inverse relationship (p<0.0001) between circulating miR-100-5p levels and both hPCSK9 levels and the magnitude of postprandial TG response **(Figure 7B).** Our observations indicate that individuals who exhibited reduced expression of miR-100-5p were characterized by elevated hPCSK9 levels and an exacerbated postprandial TG response. On the other hand, individuals with elevated miR-100-5p expression levels featured diminished circulating hPCSK9 levels and reduced postprandial hypertriglyceridemia.

**Figure 7:**
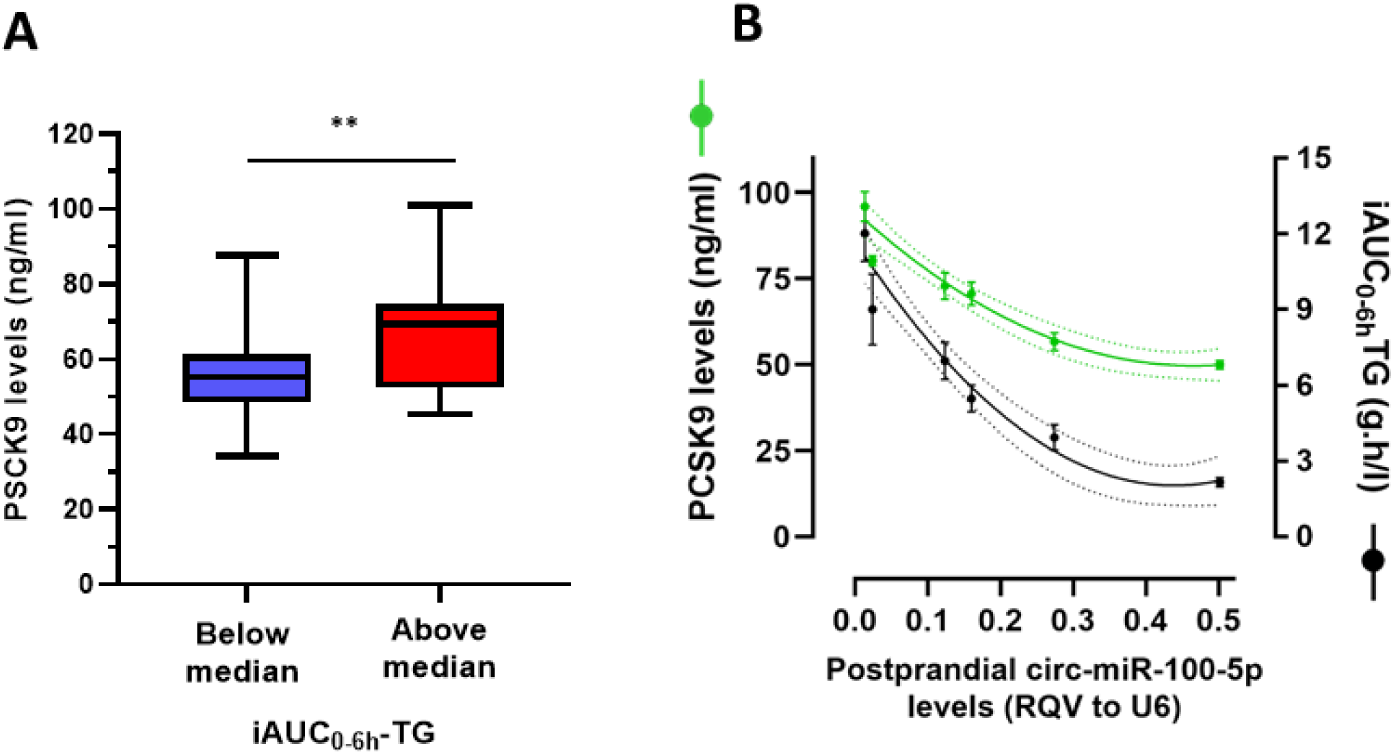
Relationship between miR-100-5p expression, postprandial TG response and PCSK9 levels. **(A)** Plasma levels of PCSK9 determined in subjects exhibited a low PP-TG response (iAUC-TG below median) or a high postprandial TG response (iAUC-TG above median). **(B)** Inverse relationship between circulating postprandial miR100-5p expression and the degree of postprandial triglyceride response expressed as netAUC_0–6h_ of triglyceride (in black) or fasting PCSK9 levels (in green). Dashed lines represent standard error. AUC indicates area under the curve; TG, triglycerides.

## DISCUSSION

We presently demonstrated that miR-100-5p regulates the magnitude of postprandial TG response by targeting hepatic PCSK9 gene expression. Our study identified a new molecular mechanism contributing to exaggerated PP-TG elevation in individuals with high risk of cardiovascular diseases. Given the established role of PCSK9 as a regulator of postprandial lipemia^27,32,33^, our data indicate that the effect of miR-100-5p on the degree of postprandial triglyceride response is mediated at least in part through its impact on hepatic PCSK9 expression.

We further report that circulating miRs can be associated with triglyceride rich lipoprotein particles. Earlier studies evaluating lipoproteins associated miRs were mostly conducted using fasting samples, from either normolipidemic or hypercholesterolemic patients displaying low circulating levels of triglycerides rich lipoproteins^13,34,35^, and none to our knowledge, by using non-fasting samples. The higher miR-100-5p content observed in postprandial TGRL particles compared to those isolated in fasting state taken together with our in vitro observations in differentiated Caco2 cells suggests that miR-100-5p may appear in the circulation concomitantly with intestinal chylomicron secretion. If confirmed, this would imply that reduced intestinal miR-100-5p expression underlies the lower postprandial circulating levels observed in subjects with an exacerbated postprandial TG response. In good agreement, enterocyte expression of miR-100-5p has been reported to play a key role in intestinal homeostasis and barrier integrity^36^. Therefore, it is likely that reduced intestinal miR-100-5p expression may disrupt the balance between proliferation and apoptosis of intestinal epithelial cells resulting in increased intestinal permeability and low-grade inflammation known to favor development of metabolic disorders, though this mechanistic chain remains to be directly tested in vivo.

The presently observed differential distribution of cir-miR-100-5p between subjects with low versus high postprandial TG responses might equally contribute to interindividual variability of the postprandial TG response through carrier-dependent mechanisms. Indeed, it has been proposed that changes in intravascular miR-carriers rather than changes in circ-miR levels per se, might exert the greatest biological impact by determining the nature of target cells and the quantity of miRNA delivered to those cells^34^. In this context is it relevant to consider that previous studies demonstrated that HDL particles isolated from individuals exhibited an elevated PP TG response displayed a reduced capacity to a deliver cholesteryl esters (CE) to hepatic cells^5^, raising the possibility that HDL-mediated hepatic delivery of miR-100-5p is similarly impaired in these individuals

The broader cardiovascular relevance of these findings is supported by converging evidence linking miR-100 to both metabolic^37,38^ and atherosclerotic^39^ diseases, ubiquitous overexpression of miR-100 in high-fat diet fed mice reduces circulating non-fasting triglyceride levels^37^, whereas pharmacological inhibition of miR100 leads to a significant increase in triglyceride levels^39^. In addition, miR-100 has been shown to have a protective role in high-fat diet-induced metabolic syndrome and liver steatosis by limiting hepatic lipid storage ^37^. In low-density lipoprotein receptor–deficient mice, the miR-100 inhibition exacerbates atherosclerotic lesion formation, leading to an unstable plaque phenotype, while miR-100 overexpression reduces plaque size and promotes a more stable plaque phenotype^39^. Mechanistically, miR-100 overexpression in endothelial cells attenuated expression of adhesion molecules and reduced monocyte infiltration into the subendothelial space^39^, consistent with an anti-atherogenic role. Importantly, while these animal data provide compelling mechanistic support, their translation to human atherosclerosis remains to be established.

In the context of postprandial metabolism specifically, endothelial cells occupy a central position: lipoprotein lipase, anchored to the luminal endothelial surface, mediates intravascular hydrolysis of TGRL, and the resulting lipolysis products are well-recognized drivers of foam cell formation and vascular inflammation^40^. In addition, whereas hepatic clearance of TGRL remnants is mediated by the LDL-R or the LRP-1, the VLDL-R primarily accounts for extra-hepatic uptake of TGRL remnant particles^41^. Interestingly, earlier studies demonstrated that overexpression of miR-100 in endothelial cells significantly decreased expression of the VLDL-R, at both mRNA and protein levels, thus reducing peripheral VLDL uptake^42^ potentially limiting peripheral TGRL uptake, an observation consistent with the predominantly extrahepatic distribution of VLDL-R under physiological conditions^41^ .

The central mechanistic axis identified in this study involves PCSK9-mediated regulation of hepatic TGRL remnant clearance. Beyond its well-characterized role in promoting lysosomal degradation of the LDL receptor (LDL-R)^43^, PCSK9 regulates additional LDL-R family members, including the lipoprotein related receptor-1 (LRP-1), the very low-density receptor (VLDL-R) or the apolipoprotein E receptor 2 (ApoER2), that collectively mediate hepatic clearance of chylomicron and VLDL remnants^44^. We therefore propose that the inverse relationship between miR-100-5p levels and the magnitude of the postprandial TG response reflects enhanced hepatic TGRL remnant uptake consequent to miR-100-5p-driven PCSK9 suppression. This model is supported by, but not yet fully proven in, human in vivo data.

Our interpretation align with earlier studies linking PCSK9 and postprandial lipemia^45^. Individuals carrying PCSK9 loss of function variants show reduced fasting and postprandial levels of triglyceride-rich lipoprotein remnants, including TG and apoB48^33^. PCSK9-deficient mice demonstrate reduced postprandial triglyceride levels following oral lipid test^32^, attributed to lowers hepatic LDL-R degradation leading to enhanced hepatic LDL-R surface expression and consequently TGRL remnant clearance. PCSK9 has also been shown to promote intestinal apoB-containing lipoprotein secretion through both LDL-R-dependent and independent mechanisms^46^, though intestinal-specific PCSK9 deletion did not alter postprandial TG levels^27^, indicating that intestinal PCSK9 is not a primary determinant of postprandial lipemia. By contrast, hepatic PCSK9 overexpression in mice produced elevated circulating PCSK9 and exacerbated postprandial lipemia, effects fully reversed by a PCSK9 monoclonal antibody directly implicating liver-derived circulating PCSK9 as the dominant regulator of postprandial lipoprotein metabolism^27^.

Consistent with this hepatocentric model, PCSK9 inhibition in healthy subjects did not significantly impact postprandial lipid metabolism^47^. However, when conducted in patients exhibiting metabolic disorders, type 2 diabetes or dysbetalipoproteinemia, interventional studies evaluating PCSK9 monoclonal antibody therapy demonstrated that PCSK9 inhibition significantly lower levels of postprandial lipids and lipoprotein remnant particles^48–51^ without altering the production rate of chylomicron by the intestine^49^. Collectively, these observations indicate that circulating PSCK9 regulates postprandial lipoprotein metabolism primarily by controlling the hepatic clearance of triglyceride-rich lipoprotein remnants particles a mechanism our findings suggest is, in turn, regulated upstream by miR-100-5p.

Finally, by studying healthy non-dyslipidemic subjects exhibiting no clinical features of metabolic disorders, we identified 13 circ-miRs with increased expression levels during the postprandial phase. Although prior studies have reported postprandial miRNA changes, the specific signatures differ considerably across studies, with only three miRNAs (miR-29a-5p, miR-16-1-5p, and miR-26b-5p) consistently identified across investigations^52,53^. This variability most likely reflects differences in study methodology, population characteristics, and the nutritional challenge employed, underscoring the need for standardized protocols in future postprandial miRNA research.

## Conclusions

Exaggerated postprandial TG response is an early marker of cardiovascular risk, preceding clinical features preceding overt metabolic dysfunction and offering a critical window for primary prevention. The present study highlighted differential postprandial circ-miRs variations according to the degree of postprandial hypertriglyceridemia, and identified miR-100-5p as a critical regulator of hepatic uptake of triglyceride rich lipoprotein particles by targeting PCSK9 in the liver a pathway of direct relevance to atherogenic dyslipidemia. Notably, reduced circulating miR-100-5p levels were associated with impaired remnant lipoprotein clearance, a mechanism increasingly implicated in accelerated atherosclerosis and residual cardiovascular risk. Prospective, large-scale studies are now warranted to establish whether circulating miR-100-5p represents a viable biomarker for the early identification of individuals at heightened risk of premature atherosclerosis, prior to the emergence of conventional metabolic risk factors.

## Data Availability

The authors declare that all supporting data are available within the article and its Supplemental Material. Raw data for individual experiments are available from the corresponding authors on reasonable request.

## Acknowledgements

The authors thank the staff of the clinical investigation center at the Pitié-Salpêtrière Hospital (CIC-1901), including nurses and technicians, for their critical assistance in implementing the HDL-PP protocol. We are also indebted to the volunteers for their cooperation.

## Authors contribution

Conceptualization: MG, WLG, PL, CLM, JML

Data acquisition: AV, AM, CN, RD, CM, SG, EF, HD

Data interpretation: MG, AV, AM, PL, WLG, OB, CM, CLM

Clinical Investigation: JS, JML

Statistical analyses: MG, AM, AV, OB

Funding acquisition: MG, WLG, PL, JML

Project administration: MG, PL, WLG

Supervision: MG, WLG, PL

Writing – original draft: AM, AV, MG

Writing – review & editing: MG, AM, AV, CM, CLM, WLG, PL, PG

## References

1. Bansal S, Buring JE, Rifai N, Mora S, Sacks FM, Ridker PM. Fasting Compared with Nonfasting Triglycerides and Risk of Cardiovascular Events in Women. J. Am. Med. Assoc. 2007, 298, 309–316, doi:10.1001/jama.298.3.309.

2. Nordestgaard BG, Benn M, Schnohr P, Tybjærg-Hansen A. Nonfasting Triglycerides and Risk of Myocardial Infarction, Ischemic Heart Disease, and Death in Men and Women. J. Am. Med. Assoc. 2007, 298, 299–308, doi:10.1001/jama.298.3.299.

3. Kolovou G, Mikhailidis DP, Kovar J, Lairon D, Nordestgaard BG, Ooi TC, Perez-Martinez P, Bilianou H, Anagnostopoulou K, Panotopoulos G. Assessment and Clinical Relevance of Non-Fasting and Postprandial Triglycerides: An Expert Panel Statement. Curr. Vasc. Pharmacol. 2011, 9, 258–270, doi:10.2174/157016111795495549.

4. Kolovou GD, Watts GF, Mikhailidis DP, Pérez-Martínez P, Mora S, Bilianou H, Panotopoulos G, Katsiki N, Ooi TC, Lopez-Miranda J, Tybjærg-Hansen A, Tentolouris N, Nordestgaard BG. Postprandial Hypertriglyceridaemia Revisited in the Era of Non-Fasting Lipid Profile Testing: A 2019 Expert Panel Statement, Main Text. Curr Vasc Pharmacol. 2019;17(5):498–514. doi: 10.2174/1570161117666190507110519.

5. Motte A, Gall J, Salem JE, Dasque E, Lebot M, Frisdal E, Galier S, Villard EF, Bouaziz-Amar E, Lacorte JM, Charbit B, Le Goff W, Lesnik P, Guerin M. Reduced Reverse Cholesterol Transport Efficacy in Healthy Men with Undesirable Postprandial Triglyceride Response. Biomolecules. 2020 May 25;10(5):810. doi: 10.3390/biom10050810.

6. Keirns BH, Sciarrillo CM, Koemel NA, Emerson SR. Fasting, Non-Fasting and Postprandial Triglycerides for Screening Cardiometabolic Risk. J. Nutr. Sci. 2021, 10, 1–14, doi:10.1017/jns.2021.73.

7. Watts GF, Ooi EMM, Chan DC. Demystifying the Management of Hypertriglyceridaemia. Nat. Rev. Cardiol. 2013, 10, 648–661, doi:10.1038/nrcardio.2013.140.

8. Nordestgaard BG. Triglyceride-Rich Lipoproteins and Atherosclerotic Cardiovascular Disease: New Insights from Epidemiology, Genetics, and Biology. Circ. Res. 2016, 118, 547–563, doi:10.1161/CIRCRESAHA.115.306249.

9. Rosenson RS, Davidson MH, Hirsh BJ, Kathiresan S, Gaudet D. Genetics and causality of triglyceride-rich lipoproteins in atherosclerotic cardiovascular disease. J Am Coll Cardiol. 2014 Dec 16;64(23):2525–40. doi: 10.1016/j.jacc.2014.09.042.

10. Ramzan F, Vickers MH, Mithen RF. Epigenetics, Microrna and Metabolic Syndrome: A Comprehensive Review. Int. J. Mol. Sci. 2021, 22, doi:10.3390/ijms22095047.

11. Parnell LD, Ordovas JM, Lai CQ. Environmental and Epigenetic Regulation of Postprandial Lipemia. Curr. Opin. Lipidol. 2018, 29, 30–35, doi:10.1097/MOL.0000000000000469.

12. Bartel DP. MicroRNAs: Genomics, Biogenesis, Mechanism, and Function. Cell 2004, 116, 281–297, doi:10.1016/S0092-8674(04)00045-5.

13. Vickers KC, Palmisano BT, Shoucri BM, Shamburek RD, Remaley AT. MicroRNAs Are Transported in Plasma and Delivered to Recipient Cells by High-Density Lipoproteins. Nat. Cell Biol. 2011, 13, 423–435, doi:10.1038/ncb2210.

14. Agbu P, Carthew RW. MicroRNA-Mediated Regulation of Glucose and Lipid Metabolism. Nat. Rev. Mol. Cell Biol. 2021, 22, 425–438, doi:10.1038/s41580-021-00354-w.

15. Desgagné V, Bouchard L, Guérin R. MicroRNAs in Lipoprotein and Lipid Metabolism: From Biological Function to Clinical Application. Clin. Chem. Lab. Med. 2017, 55, 667–686, doi:10.1515/cclm-2016-0575.

16. Fernández-Tussy P, Ruz-Maldonado I, Fernández-Hernando C. MicroRNAs and Circular RNAs in Lipoprotein Metabolism. Curr Atheroscler Rep. 2021 May 10;23(7):33. doi: 10.1007/s11883-021-00934-3.

17. Sun C, Alkhoury K, Wang YI, Foster GA, Radecke CE, Tam K, Edwards CM, Facciotti MT, Armstrong EJ, Knowlton AA, Newman JW, Passerini AG, Simon SI. IRF-1 and miRNA126 modulate VCAM-1 expression in response to a high-fat meal. Circ Res. 2012 Sep 28;111(8):1054–64. doi: 10.1161/CIRCRESAHA.112.270314.

18. Zhu T, Corraze G, Plagnes-Juan E, Quillet E, Dupont-Nivet M, Skiba-Cassy S. Regulation of Genes Related to Cholesterol Metabolism in Rainbow Trout ( Oncorhynchus Mykiss ) Fed a Plant-Based Diet . Am. J. Physiol. Integr. Comp. Physiol. 2017, 314, R58–R70, doi:10.1152/ajpregu.00179.2017.

19. Mennigen JA, Panserat S, Larquier M, Plagnes-Juan E, Medale F, Seiliez I, Skiba-Cassy S. Postprandial Regulation of Hepatic MicroRNAs Predicted to Target the Insulin Pathway in Rainbow Trout. PLoS One 2012, 7, doi:10.1371/journal.pone.0038604.

20. Mantilla-Escalante DC, López de Las Hazas MC, Gil-Zamorano J, Del Pozo-Acebo L, Crespo MC, Martín-Hernández R, Del Saz A, Tomé-Carneiro J, Cardona F, Cornejo-Pareja I, García-Ruiz A, Briand O, Lasunción MA, Visioli F, Dávalos A. Postprandial Circulating miRNAs in Response to a Dietary Fat Challenge. Nutrients. 2019 Jun 13;11(6):1326. doi: 10.3390/nu11061326.

21. Lopez S, Bermudez B, Montserrat-de la Paz S, Abia R, Muriana FJG. A MicroRNA Expression Signature of the Postprandial State in Response to a High-Saturated-Fat Challenge. J. Nutr. Biochem. 2018, 57, 45–55, doi:10.1016/j.jnutbio.2018.03.010.

22. Talbot CPJ, Mensink RP, Smolders L, Bakeroot V, Plat J. Theobromine Does Not Affect Fasting and Postprandial HDL Cholesterol Efflux Capacity, While It Decreases Fasting MiR-92a Levels in Humans. Mol. Nutr. Food Res. 2018, 62, 1–8, doi:10.1002/mnfr.201800027.

23. Ramzan F, D’Souza RF, Durainayagam BR, Milan AM, Roy NC, Kruger MC, Henry CJ, Mitchell CJ, Cameron-Smith D. Inflexibility of the Plasma MiRNA Response Following a High-Carbohydrate Meal in Overweight Insulin-Resistant Women. Genes Nutr. 2020, 15, 1–12, doi:10.1186/s12263-020-0660-8.

24. Lassel TS, Guérin M, Auboiron S, Chapman MJ, Guy-Grand B. Preferential cholesteryl ester acceptors among triglyceride-rich lipoproteins during alimentary lipemia in normolipidemic subjects. Arterioscler Thromb Vasc Biol. 1998 Jan;18(1):65–74. doi: 10.1161/01.atv.18.1.65.

25. Nauli AM, Sun Y, Whittimore JD, Atyia S, Krishnaswamy G, Nauli SM. Chylomicrons produced by Caco-2 cells contained ApoB-48 with diameter of 80-200 nm. Physiol Rep. 2014 Jun 6;2(6):e12018. doi: 10.14814/phy2.12018.

26. Luchoomun J, Hussain MM. Assembly and secretion of chylomicrons by differentiated Caco-2 cells. Nascent triglycerides and preformed phospholipids are preferentially used for lipoprotein assembly. J Biol Chem. 1999 Jul 9;274(28):19565–72. doi: 10.1074/jbc.274.28.19565.

27. Garçon D, Moreau F, Ayer A, Dijk W, Prieur X, Arnaud L, Roubtsova A, Seidah N, Prat A, Cariou B, Le May C. Circulating Rather Than Intestinal PCSK9 (Proprotein Convertase Subtilisin Kexin Type 9) Regulates Postprandial Lipemia in Mice. Arterioscler Thromb Vasc Biol. 2020 Sep;40(9):2084–2094. doi: 10.1161/ATVBAHA.120.314194.

28. Bruckert, E.; Le Goff, W. Physiologie Du Métabolisme Des Lipoprotéines. Med. des Mal. Metab. 2018, 12, 50–61, doi:10.1016/S1957-2557(18)30009-9.

29. Zhang BH, Yin F, Qiao YN, Guo SD. Triglyceride and Triglyceride-Rich Lipoproteins in Atherosclerosis. Front Mol Biosci. 2022 May 25;9:909151. doi: 10.3389/fmolb.2022.909151.

30. Chen Y, Wang X. MiRDB: An Online Database for Prediction of Functional MicroRNA Targets. Nucleic Acids Res. 2020, 48, D127–D131, doi:10.1093/nar/gkz757.

31. Chang L, Zhou G, Soufan O, Xia J. MiRNet 2.0: Network-Based Visual Analytics for MiRNA Functional Analysis and Systems Biology. Nucleic Acids Res. 2020, 48, W244–W251, doi:10.1093/nar/gkaa467.

32. Le May C, Kourimate S, Langhi C, Chétiveaux M, Jarry A, Comera C, Collet X, Kuipers F, Krempf M, Cariou B, Costet P. Proprotein convertase subtilisin kexin type 9 null mice are protected from postprandial triglyceridemia. Arterioscler Thromb Vasc Biol. 2009 May;29(5):684–90. doi: 10.1161/ATVBAHA.108.181586.

33. Ooi TC, Krysa JA, Chaker S, Abujrad H, Mayne J, Henry K, Cousins M, Raymond A, Favreau C, Taljaard M, Chrétien M, Mbikay M, Proctor SD, Vine DF. The Effect of PCSK9 Loss-of-Function Variants on the Postprandial Lipid and ApoB-Lipoprotein Response. J Clin Endocrinol Metab. 2017 Sep 1;102(9):3452–3460. doi: 10.1210/jc.2017-00684.

34. Desgagné V, Guérin R, Guay SP, Corbin F, Couture P, Lamarche B, Bouchard L. Changes in High-Density Lipoprotein-Carried MiRNA Contribution to the Plasmatic Pool after Consumption of Dietary Trans Fat in Healthy Men. Futur. Med. 2017, 9, 669–688, doi:10.2217/epi-2016-0177.

35. Wagner, J.; Riwanto, M.; Besler, C.; Knau, A.; Fichtlscherer, S.; Röxe, T.; Zeiher, A.M.; Landmesser, U.; Dimmeler, S. Characterization of Levels and Cellular Transfer of Circulating Lipoprotein-Bound MicroRNAs. Arterioscler. Thromb. Vasc. Biol. 2013, 33, 1392–1400, doi:10.1161/ATVBAHA.112.300741.

36. Zou, L.; Xiong, X.; Yang, H.; Wang, K.; Zhou, J.; Lv, D.; Yin, Y. Identification of MicroRNA Transcriptome Reveals That MiR-100 Is Involved in the Renewal of Porcine Intestinal Epithelial Cells. Sci. China Life Sci. 2019, 62, 816–828, doi:10.1007/s11427-018-9338-9.

37. Smolka C, Schlösser D, Hohnloser C, Bemtgen X, Jänich C, Schneider L, Martin J, Pfeifer D, Moser M, Hasselblatt P, Bode C, Grundmann S, Pankratz F. MiR-100 overexpression attenuates high fat diet induced weight gain, liver steatosis, hypertriglyceridemia and development of metabolic syndrome in mice. Mol Med. 2021 Sep 6;27(1):101. doi: 10.1186/s10020-021-00364-6.

38. Pek, S.L.T., Sum, C.F., Lin, M.X., Cheng, A.K.S., Wong, M.T.K., Lim, S.C., Tavintharan, S. Circulating and Visceral Adipose MiR-100 Is down-Regulated in Patients with Obesity and Type 2 Diabetes. Mol. Cell. Endocrinol. 2016, 427, 112–123, doi:10.1016/j.mce.2016.03.010.

39. Pankratz F, Hohnloser C, Bemtgen X, Jaenich C, Kreuzaler S, Hoefer I, Pasterkamp G, Mastroianni J, Zeiser R, Smolka C, Schneider L, Martin J, Juschkat M, Helbing T, Moser M, Bode C, Grundmann S. MicroRNA-100 Suppresses Chronic Vascular Inflammation by Stimulation of Endothelial Autophagy. Circ Res. 2018 Feb 2,122(3):417-432. doi: 10.1161/CIRCRESAHA.117.311428.

40. Schwartz EA, Reaven PD. Lipolysis of triglyceride-rich lipoproteins, vascular inflammation, and atherosclerosis. Biochim Biophys Acta. 2012 May,1821(5):858–66. doi: 10.1016/j.bbalip.2011.09.021.

41. Takahashi S, Sakai J, Fujino T, Hattori H, Zenimaru Y, Suzuki J, Miyamori I, Yamamoto TT. The very low-density lipoprotein (VLDL) receptor: characterization and functions as a peripheral lipoprotein receptor. J Atheroscler Thromb. 2004,11(4):200–8. doi: 10.5551/jat.11.200.

42. Fang X, Fang L, Liu A, Wang X, Zhao B, Wang N. Activation of PPAR-δ Induces MicroRNA-100 and Decreases the Uptake of Very Low-Density Lipoprotein in Endothelial Cells. Br. J. Pharmacol. 2015, 172, 3728–3736, doi:10.1111/bph.13160.

43. Xia XD, Peng ZS, Gu HM, Wang M, Wang GQ, Zhang DW. Regulation of PCSK9 Expression and Function: Mechanisms and Therapeutic Implications. Front Cardiovasc Med. 2021 Oct 15,8:764038. doi: 10.3389/fcvm.2021.764038.

44. Poirier S, Mayer G, Benjannet S, Bergeron E, Marcinkiewicz J, Nassoury N, Mayer H, Nimpf J, Prat A, Seidah NG. The proprotein convertase PCSK9 induces the degradation of low density lipoprotein receptor (LDLR) and its closest family members VLDLR and ApoER2. J Biol Chem. 2008 Jan 25,283(4):2363–72. doi: 10.1074/jbc.M708098200.

45. Druce I, Abujrad H, Ooi TC. PCSK9 and triglyceride-rich lipoprotein metabolism. J Biomed Res. 2015 Nov,29(6):429–436. doi: 10.7555/JBR.29.20150052.

46. Rashid S, Tavori H, Brown PE, Linton MF, He J, Giunzioni I, Fazio S. Proprotein convertase subtilisin kexin type 9 promotes intestinal overproduction of triglyceride-rich apolipoprotein B lipoproteins through both low-density lipoprotein receptor-dependent and - independent mechanisms. Circulation. 2014 Jul 29,130(5):431–41. doi: 10.1161/CIRCULATIONAHA.113.006720. Epub 2014 May 23. Erratum in: Circulation. 2015 May 5,131(18):e429. doi: 10.1161/CIR.0000000000000210.

47. Reyes-Soffer G, Pavlyha M, Ngai C, Thomas T, Holleran S, Ramakrishnan R, Karmally W, Nandakumar R, Fontanez N, Obunike J, Marcovina SM, Lichtenstein AH, Matthan NR, Matta J, Maroccia M, Becue F, Poitiers F, Swanson B, Cowan L, Sasiela WJ, Surks HK, Ginsberg HN. Effects of PCSK9 Inhibition With Alirocumab on Lipoprotein Metabolism in Healthy Humans. Circulation. 2017 Jan 24,135(4):352–362. doi: 10.1161/CIRCULATIONAHA.116.025253. Epub 2016 Dec 16.

48. Burggraaf B, Pouw NMC, Arroyo SF, van Vark-van der Zee LC, van de Geijn GM, Birnie E, Huisbrink J, van der Zwan EM, Mulder MT, Rensen PCN, de Herder WW, Cabezas MC. A placebo-controlled proof-of-concept study of alirocumab on postprandial lipids and vascular elasticity in insulin-treated patients with type 2 diabetes mellitus. Diabetes Obes Metab. 2020 May,22(5):807–816. doi: 10.1111/dom.13960. Epub 2020 Jan 27.

49. Taskinen MR, Björnson E, Andersson L, Kahri J, Porthan K, Matikainen N, Söderlund S, Pietiläinen K, Hakkarainen A, Lundbom N, Nilsson R, Ståhlman M, Adiels M, Parini P, Packard C, Borén J. Impact of proprotein convertase subtilisin/kexin type 9 inhibition with evolocumab on the postprandial responses of triglyceride-rich lipoproteins in type II diabetic subjects. J Clin Lipidol. 2020 Jan-Feb,14(1):77–87. doi: 10.1016/j.jacl.2019.12.003.

50. Heidemann BE, Koopal C, Roeters van Lennep JE, Stroes ESG, Riksen NP, Mulder MT, - van der Zee LCVV, Blackhurst DM, Marais AD, Visseren FLJ. Effect of evolocumab on fasting and post fat load lipids and lipoproteins in familial dysbetalipoproteinemia. J Clin Lipidol. 2023 Jan-Feb,17(1):112–123. doi: 10.1016/j.jacl.2022.10.006.

51. Heidemann BE, Marais AD, Mulder MT, Visseren FLJ, Roeters van Lennep JE, Stroes ESG, Riksen NP, van Vark-van der Zee LC, Blackhurst DM, Koopal C. Composition and distribution of lipoproteins after evolocumab in familial dysbetalipoproteinemia: A randomized controlled trial. J Clin Lipidol. 2023 Sep-Oct,17(5):666–676. doi: 10.1016/j.jacl.2023.07.004.

52. Daimiel L, Micó V, Valls RM, Pedret A, Motilva MJ, Rubió L, Fitó M, Farrás M, Covas MI, Solá R, Ordovás JM. Impact of Phenol-Enriched Virgin Olive Oils on the Postprandial Levels of Circulating microRNAs Related to Cardiovascular Disease. Mol Nutr Food Res. 2020 Aug;64(15):e2000049. doi: 10.1002/mnfr.202000049.

53. Quintanilha BJ, Pinto Ferreira LR, Ferreira FM, Neto EC, Sampaio GR, Rogero MM. Circulating Plasma MicroRNAs Dysregulation and Metabolic Endotoxemia Induced by a High-Fat High-Saturated Diet. Clin. Nutr. 2020, 39, 554–562, doi:10.1016/j.clnu.2019.02.042.

