## Supplemental data for "miR-100-5p modulates postprandial triglyceride response by targeting PCSK9"

**Members of the RESIST-PP Consortium:**

Stephan C. BISCHOFF[1], Mathias CHAMAILLARD[2], Maryse GUERIN[3], Lucy HEZINGER[1], Pukar KC[2], Thomas KUFER[1], Martin LARSEN[4], Philippe LESNIK[3], Juan José MARTINEZ-GARCIA[5], Carolina NEVES[3], Maharajah PONNAIAH[6], Pablo PELEGRIN[5], Joe-Elie SALEM[7].

**Affiliations**

[1] Institute of Nutritional Medicine, Department of Immunology, University of Hohenheim, 70593 Stuttgart, Germany.

[2]Inserm, University of Lille, U1003, F-59000, Lille, France.

[3]Sorbonne Université, Inserm, UMR_S1166, Research Institute of Cardiovascular Disease, Metabolism and Nutrition, Faculté de Santé - Hôpital Pitié-Salpêtrière, Paris, France

[4]Sorbonne Université, Inserm U1135, CNRS EMR 8255, Centre d'Immunologie et des Maladies Infectieuses (CIMI-Paris), Paris, France.

[5]Department of Biochemistry and Molecular Biology B and Immunology, Faculty of Medicine, University of Murcia, 30120, Murcia, Spain.

[6]Foundation for Innovation in Cardiometabolism and Nutrition (IHU ICAN, ICAN OMICS and ICAN I/O), F-75013 Paris, France

[7]Centre d’Investigation Clinique Paris-Est CIC-1901, Hôpital de la Pitié-Salpêtrière, AP-HP, Paris, France

**Supplemental Table S1. Detected circulating miRs with reliable results for healthy non-dyslipidemic subjects included in TaqMan Low-density Arrays (TLDAs) and differentially regulated according to the degree of postprandial TG response.**

| Identified circulating miRs | Ct values (cycles) |
|---|---|
| *Human TaqMan miRNA arrays Card A v3.0:*<br>hsa-miR-223-3p; hsa-miR-24-3p; hsa-miR-19b-3p; hsa-miR-484; hsa-miR-191-5p; hsa-miR-146a-5p; hsa-miR-106a-5p; hsa-miR-126-3p; hsa-miR-17-5p hsa-miR-20a-5p; **hsa-miR-16-5p**; hsa-miR-92a-3p<br><br>*Human TaqMan miRNA arrays Card B v3.0:*<br>None | 15-20 |
| *Human TaqMan miRNA arrays Card A v3.0:*<br>hsa-miR-222-3p; hsa-miR-320a-3p; hsa-miR-221-3p; hsa-miR-30c-5p; hsa-miR-21-5p; hsa-miR-145-5p; hsa-miR-342-3p; hsa-let-7b-5p; hsa-miR-26a-5p; hsa-miR-30b-5p; hsa-miR-328-3p; hsa-let-7e-5p; hsa-miR-197-3p; hsa-miR-199a-3p; hsa-miR-186-5p; hsa-let-7a-5p; hsa-miR-425-5p; hsa-miR-142-3p; hsa-miR-146b-5p; hsa-miR-486-5p; hsa-miR-20b-5p; hsa-let-7d-5p; hsa-miR-15b-5p; hsa-miR-19a-3p; hsa-miR-103a-3p; hsa-miR-331-3p; hsa-miR-150-5p; hsa-miR-335-5p; hsa-let-7g-5p; hsa-miR-28-3p; hsa-miR-106b-5p; **hsa-miR-26b-5p**; hsa-miR-574-3p; hsa-miR-27a-3p; hsa-miR-25-3p; hsa-miR-18a-5p; hsa-let-7f-5p; hsa-miR-130a-3p; hsa-miR-99b-5p; hsa-miR-18b-5p; hsa-miR-652-3p; hsa-miR-744-5p; hsa-miR-23a-3p; hsa-miR-98-5p; hsa-miR-301a-3p; hsa-miR-339-5p; hsa-miR-423-5p; hsa-miR-374-5p; hsa-miR-195-5p; hsa-miR-382-5p.<br><br>*Human TaqMan miRNA arrays Card B v3.0:*<br>hsa-miR-1274B; hsa-miR-126-5p; hsa-miR-151a-3p; hsa-miR-30a-5p; hsa-miR-766-3p; hsa-miR-409-3p; hsa-miR-720; hsa-miR-30d-5p; hsa-miR-151a-5p; hsa-miR-625-3p; hsa-miR-30e-3p. | 20-25 |
| *Human TaqMan miRNA arrays Card A v3.0:*<br>hsa-miR-133a-3p; hsa-miR-130b-3p; hsa-miR-454-3p; hsa-miR-532-5p; hsa-miR-139-5p; hsa-miR-122-5p; hsa-miR-339-3p; hsa-miR-27b-3p; hsa-miR-181a-5p; hsa-miR-324-5p; hsa-miR-28-5p; hsa-miR-125a-5p; hsa-miR-29a-3p; hsa-miR-345-5p; hsa-miR-132-3p; hsa-miR-224; hsa-miR-185-5p; hsa-miR-376c-3p; hsa-miR-210-3p; hsa-miR-152-3p; hsa-miR-127-3p; hsa-miR-148a-3p; hsa-miR-15a-5p; hsa-miR-128-3p; hsa-miR-485-3p; hsa-miR-330-3p; hsa-miR-660-5p; hsa-miR-340-5p; hsa-miR-155; hsa-miR-501-5p; hsa-miR-200c-3p; hsa-miR-361-5p; hsa-miR-143-3p; hsa-miR-192-5p; hsa-miR-148b-3p; hsa-miR-142-5p; hsa-miR-590-5p; hsa-miR-370-3p; hsa-miR-539-5p; hsa-miR-532-3p hsa-miR-296-5p; hsa-miR-324-3p; hsa-miR-375-3p; hsa-miR-433-3p; hsa-miR-133b; hsa-miR-376a-3p; hsa-miR-494-3p; hsa-miR-885-5p, U6 snRNA**; hsa-miR-140-3p**; hsa-miR-23b; hsa-miR-598-3p; hsa-let-7c-5p; hsa-miR-671-3p; hsa-miR-625-5p; hsa-miR-215-5p; hsa-miR-29c-3p; hsa-miR-486-3p; hsa-miR-505-3p; hsa-miR-200b-3p; hsa-miR-323a-3p; hsa-miR-483-5p; hsa-miR-200a-3p; hsa-miR-193b-3p; hsa-miR-365a-3p; hsa-miR-10a-5p; hsa-miR-101-3p; hsa-miR-410-3p; hsa-miR-34a-5p; hsa-miR-886-5p; **hsa-miR-1-3p**; hsa-miR-193a-5p; hsa-miR-485-5p; hsa-miR-204-5p; hsa-miR-301b-3p; hsa-miR-99a-5p; hsa-miR-211-5p; hsa-miR-487b-3p; hsa-miR-874-3p; hsa-miR-194-5p; hsa-miR-409-5p; hsa-miR-501-3p; hsa-miR-628-5p; hsa-miR-597-5p; hsa-miR-363-3p; hsa-miR-331-5p; hsa-miR-579-3p; hsa-miR-181c-5p; hsa-miR-199a-5p; **hsa-miR-100-5p**; hsa-miR-346; hsa-miR-342-5p<br><br>*Human TaqMan miRNA arrays Card B v3.0:*<br>hsa-miR-664a-3p; hsa-miR-223-5p; hsa-miR-22-5p; hsa-miR-93-3p; hsa-miR-1066-3p; hsa-miR-432-5p; hsa-miR-483-3p; hsa-miR-320b; hsa-miR-1298-5p; hsa-miR-425-3p; hsa-miR-584-5p; hsa-miR-30a-3p; hsa-miR-1274A; U6 snRNA; hsa-miR-340-3p; hsa- | 25-30 |

| | |
|---|---|
| miR-335-3p; hsa-miR-378a-3p; hsa-miR-130b-5p; hsa-miR-26b-3p; hsa-miR-1233-3p; hsa-miR-1301-3p; hsa-miR-769-5p; hsa-miR-543; hsa-miR-1825; hsa-miR-505-5p; hsa-miR-15b-3p; hsa-miR-1260a; hsa-miR-1249-3p; hsa-miR-628-3p; hsa-miR-361-3p; hsa-miR-181c-3p; hsa-miR-1296-5p; hsa-miR-942-5p; hsa-miR-144-5p; hsa-miR-181a-2-3p; hsa-miR-191-3p; hsa-miR-15a-3p; hsa-miR-26a-1-3p; hsa-miR-590-3p; hsa-miR-1290; hsa-miR-1271-5p; hsa-miR-338-5p; hsa-miR-589-3p; hsa-miR-99b-3p; hsa-miR-10b-3p; hsa-miR-571; hsa-miR-424-3p. | |
| *Human TaqMan miRNA arrays Card A v3.0:*<br>hsa-miR-125a-3p; hsa-miR-326; hsa-miR-411-5p; hsa-miR-636; hsa-miR-452-5p; hsa-miR-214-3p; hsa-miR-431-5p; hsa-miR-629-5p; hsa-miR-500a-5p; hsa-miR-576-3p; hsa-miR-337-5p; hsa-miR-196b-5p; hsa-miR-518d-3p; hsa-miR-655-3p; hsa-miR-138-5p; hsa-miR-502-3p; hsa-miR-758-3p; hsa-miR-199b-5p; hsa-miR-487a-3p; hsa-miR-362-5p; hsa-miR-654-5p; hsa-miR-582-3p; hsa-miR-493-3p; hsa-miR-889-3p; hsa-miR-212-3p; **hsa-miR-9-5p**; hsa-miR-146b-3p; hsa-miR-654-3p; hsa-miR-545-3p; hsa-miR-29b-3p; hsa-miR-141-3p; hsa-miR-338-3p; hsa-miR-369-5p; hsa-miR-329-3p; hsa-miR-519a-3p; hsa-miR-886-3p; hsa-miR-362-3p; hsa-miR-616-3p.<br><br>*Human TaqMan miRNA arrays Card B v3.0:*<br>hsa-miR-144-3p; hsa-miR-183-3p; hsa-miR-629-3p; hsa-let-7b-3p; hsa-miR-19b-1-5p; hsa-miR-18a-3p; hsa-miR-181a-3p; hsa-miR-21-3p; hsa-miR-9-3p; hsa-miR-33a-5p; hsa-miR-454-5p; hsa-miR-222-5p; hsa-miR-1227-3p; hsa-let-7f-1-3p; hsa-miR-145-3p; hsa-miR-24-2-5p; hsa-miR-337-3p; hsa-miR-190b; hsa-miR-661; hsa-miR-1208; hsa-miR-25-5p; hsa-miR-148b-5p; hsa-miR-20a-3p; hsa-miR-1255b-5p; hsa-miR-136-3p; hsa-miR-548j-5p; hsa-miR-185-3p; hsa-miR-1180-3p; hsa-miR-1285-3p; **hsa-miR-941**; **hsa-miR-744-3p;** hsa-miR-33a-3p; hsa-miR-638; hsa-miR-27b-5p; hsa-miR-550a-3p; hsa-miR-497-5p; hsa-miR-616-5p; **hsa-miR-29b-2-5p**; hsa-miR-1183; hsa-miR-17-3p; hsa-miR-30d-3p; **hsa-miR-154-3p;** hsa-miR-1197; hsa-miR-605-5p; hsa-let-7a-3p; hsa-miR-656-3p; hsa-miR-34a-3p; hsa-miR-1247-5p; hsa-miR-23b-5p; **hsa-miR-29a-5p;** hsa-miR-148a-5p; **hsa-miR-16-1-3p.** | 30-35 |

miRNA detection is represented as normalized raw cycle threshold (Ct) values from RT-PCR analysis.
Circulating miRs associated with postprandial TG response are indicated in bold.

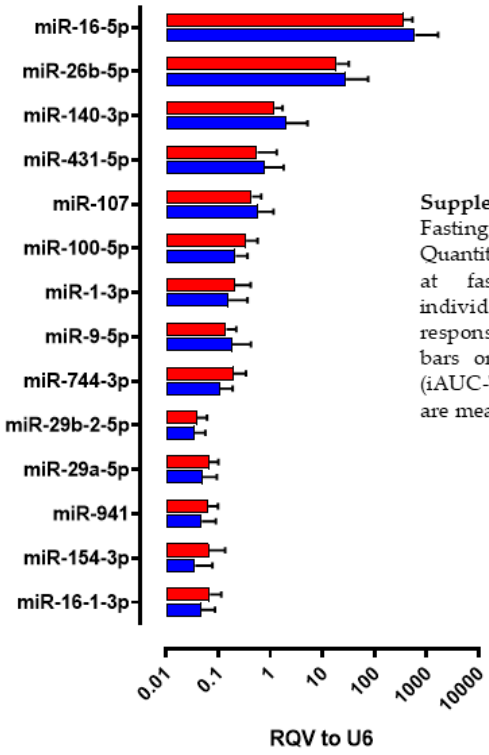


**Supplemental Figure S1.**
Fasting circulating miR levels. Relative Quantitative Value (RQV) to U6 of circ-miRs at fasting state in non-dyslipidemic individuals exhibited a low postprandial TG response (iAUC-TG below median, blue bars or a high postprandial TG response (iAUC-TG above median, red bars). Values are mean ±SD.

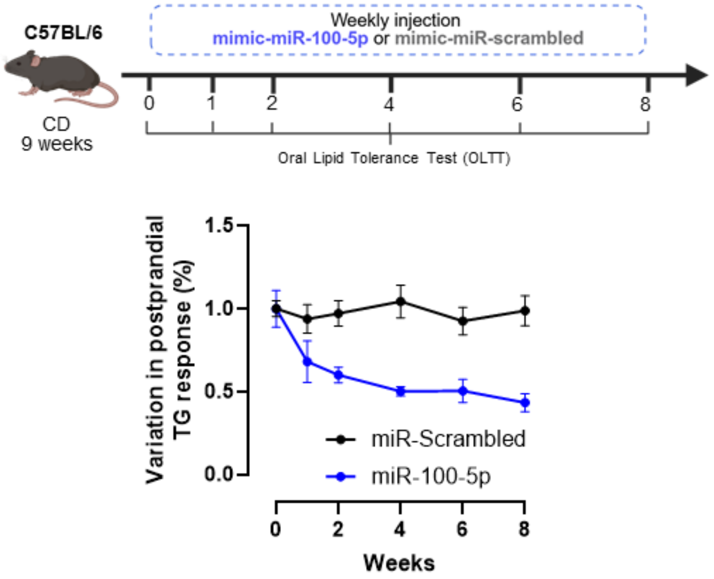


**Supplemental Figure S2.**
Variation in postprandial TG response of mice after 1, 2, 4, 6 and 8 weeks of weekly injection of mimic miR-Scrambled (n=10) or mimic miR-100-5p (n=10).
